# Convergence of a common solution to broad ebolavirus neutralization by glycan cap directed human antibodies

**DOI:** 10.1101/2020.10.14.340026

**Authors:** Charles D. Murin, Pavlo Gilchuk, Philipp A. Ilinykh, Kai Huang, Natalia Kuzmina, Xiaoli Shen, Jessica F. Bruhn, Aubrey L. Bryan, Edgar Davidson, Benjamin J. Doranz, Lauren E. Williamson, Jeffrey Copps, Tanwee Alkutkar, Andrew I. Flyak, Alexander Bukreyev, James E. Crowe, Andrew B. Ward

## Abstract

Antibodies that target the glycan cap epitope on ebolavirus glycoprotein (GP) are common in the adaptive response of survivors. A subset is known to be broadly neutralizing, but the details of their epitopes and basis for neutralization is not well-understood. Here we present cryo-electron microscopy (cryo-EM) structures of several glycan cap antibodies that variably synergize with GP base-binding antibodies. These structures describe a conserved site of vulnerability that anchors the mucin-like domains (MLD) to the glycan cap, which we name the MLD-anchor and cradle. Antibodies that bind to the MLD-cradle share common features, including the use of *IGHV1-69* and *IGHJ6* germline genes, which exploit hydrophobic residues and form beta-hairpin structures to mimic the MLD-anchor, disrupt MLD attachment, destabilize GP quaternary structure and block cleavage events required for receptor binding. Our results collectively provide a molecular basis for ebolavirus neutralization by broadly reactive glycan cap antibodies.

## Introduction

There is mounting evidence that protection from filoviral infection can be achieved through the use of monoclonal antibodies (mAbs) that target the GP surface (Bornholdt et al., 2019; Brannan et al., 2019; Mire et al., 2017; Qiu et al., 2014; Saphire et al., 2018b). Several structures of antigen-antibody complexes in recent years indicate that antibodies can access nearly any region on the surface of GP (Flyak et al., 2015; Flyak et al., 2018; Gilchuk et al., 2018; Milligan et al., 2019; Murin, 2018; Murin et al., 2019; Pallesen et al., 2016; Pascal et al., 2018; Saphire et al., 2018a; Wec et al., 2017; West et al., 2018; Zhao et al., 2017). Such antibodies have utility as post-exposure therapeutics when used in combination, such as the tri-mAb cocktail REGN-EB3, which demonstrated high efficacy in animal models (Pascal et al., 2018) and in a clinical trial carried out during a recent Ebola virus (EBOV) outbreak (Mulangu et al., 2019). REGN-EB3 is only effective against EBOV, however a pan-ebolavirus therapeutic that recognizes multiple ebolaviruses that cause severe disease in humans and major outbreaks, including Bundibugyo virus (BDBV) and Sudan virus (SUDV), would be ideal given the unpredictability of ebolavirus outbreaks.

Cross-reactive antibodies often target regions of conserved sequence vital to viral function, such as the receptor binding site (RBS) (Flyak et al., 2015; Hashiguchi et al., 2015; Howell et al., 2016; King et al., 2018), the internal fusion loop (IFL) (Milligan et al., 2019; Murin, 2018; West et al., 2018; Zhao et al., 2017), the base of GP (Gilchuk et al., 2018; Misasi et al., 2016) and the heptad repeat 2 (HR2) region (Bornholdt et al., 2016b; Flyak et al., 2018). Less conserved regions, such as the glycan cap and MLD, also can be targeted by protective antibodies and typically represent the largest antibody responses found in survivors; however, such antibodies are usually weakly or non-neutralizing and species-specific (Murin et al., 2014; Zeitlin et al., 2011). For example, the antibody 13C6, which was included in the antibody cocktail ZMapp^TM^, targets the glycan cap, but is low in potency for viral neutralization and is thought to instead provide protection by facilitating a superior cellular response (Murin et al., 2014; Pallesen et al., 2016). Furthermore, the glycan cap/head epitope in the trimeric membrane form of GP is also partially present on sGP, the soluble dimer of GP that is secreted in abundance during natural infection (Cook and Lee, 2013; de La Vega et al., 2015; Pallesen et al., 2016). Finally, GP is massively remodeled during endosomal entry in processes mediated by host proteases, during which the glycan cap and MLD are removed (Bornholdt et al., 2016a; Lee and Saphire, 2009). Nevertheless, several antibodies have been identified that bind within the glycan cap and potently neutralize EBOV, BDBV and SUDV (Bornholdt et al., 2016b; Flyak et al., 2016; Gilchuk et al., 2020; Misasi et al., 2016; Murin et al., 2014; Pascal et al., 2018; Saphire et al., 2018a). The mechanistic basis for this activity, however, is not well-explored.

We previously characterized pan-ebolavirus neutralizing mAbs isolated from a survivor cohort of the EBOV 2013-2016 outbreak (Gilchuk et al., 2018; Gilchuk et al., 2020). Several antibodies that recognize the glycan cap revealed synergistic activity for the GP binding and virus neutralization when paired with GP base-binding antibodies. One such pair, EBOV-548 and EBOV-520, provided superior protection in animal models when compared with treatment by a single antibody. Structural evaluation revealed that EBOV-520 recognized the 3^10^ pocket that is partially shielded by the β17-β18 loop in uncleaved GP. EBOV-548, which binds to the glycan cap, removed this steric hindrance by dislodging and mimicking the β18-β18′ hairpin obscuring the 3^10^ pocket. These data revealed a structural mechanism for synergy mediated by a glycan cap-directed antibody.

We sought to determine if glycan cap antibodies from other survivors also use similar mechanisms of protection and synergy as the EBOV-548/EBOV-520 combination (Bramble et al., 2018; Flyak et al., 2015; Flyak et al., 2018; Flyak et al., 2016; Gilchuk et al., 2018). This collection of antibodies, including two mAbs from a newly described survivor cohort, were tested for their ability to enhance the activity of the GP base-region-binding broadly neutralizing antibodies EBOV-520 and EBOV-515 (Gilchuk et al., 2020). Additionally, we observed and quantified antibody-induced GP trimer instability. Subsequent analysis by cryo-EM revealed a conserved structural motif, similar to that found in EBOV-548, wherein a complementarity determining region (CDR) exhibited molecular mimicry of the β18-β18′ hairpin in GP. Finally, we also quantified the ability of glycan cap antibodies to block GP cleavage events necessary for receptor binding site exposure. Our data collectively provide evidence for a mechanism behind the activity of broadly neutralizing and synergistic glycan cap antibodies to ebolaviruses and suggest a rational strategy for the design of therapeutic antibody cocktails.

## Results

### Glycan cap antibody synergy is a common feature and is associated with GP instability

We previously described an assay to determine antibody binding synergy of pairs of antibodies for glycan cap antibody-based synergy of the base-region-binding antibodies EBOV-515 and EBOV-520 (Gilchuk et al., 2020). Here, we extended this assay to glycan cap antibodies from other survivor cohorts. We chose previously isolated antibodies based on similar properties to the synergistic glycan cap mAb EBOV-548, including: 1) synergy with EBOV-520 and/or EBOV-515, 2) broad reactivity and neutralization, 3) long CDRH3 loops, 4) cross-reactivity with sGP and/or 5) protection *in vivo*. Based on these criteria, we chose the following antibodies: BDBV-43, BDBV-329, BDBV-289, EBOV-442, EBOV-437 and EBOV-237 (Flyak et al., 2016; Gilchuk et al., 2018; Qiu et al., 2012; Williamson et al., 2019; Wilson et al., 2000). EBOV-548, 13C6, and an unrelated human mAb directed to dengue virus (DENV) envelope protein 2D22, were included for comparative purposes and as controls (**Table 1**). In addition, we also tested two new antibodies, EBOV-293 and EBOV-296, which we isolated from an individual treated for EBOV infection in the United States (**Table 1**, also see *Materials and Methods*). Ten characterized glycan cap antibodies potently bound to the sGP as judged by the half-maximal effective (EC_50_) concentrations, and revealed diverse GP reactivity and virus neutralization profiles, and diverse protective efficacy in EBOV challenge mouse model (**Table 1**, **Fig. S1A**). In addition, epitope mapping by alanine scanning mutagenesis library analysis identified key contact residues for each antibody (**Table 1**; **Fig. S1B**). Furthermore, several of these antibodies have exceptionally long CDRH3 loops, such as the 33 amino acid loop of BDBV-329 (**Table S1**).

**Table 1.** Summary of binding and functional activities of characterized antibodies. Characteristics for previously described antibodies were included for comparative purposes. *Determined in this study; ** prophylaxis efficacy; ***incomplete neutralization at highest tested Ab concentration (0.2 mg/mL); “>” indicates activity was not detected at highest tested Ab concentration; ND – not determined. See also Figure S1 and Table S1.

| Antibody | GP binding by ELISA, EC<sub>50</sub><br>(ng/mL) |  |  |  | Virus neutralization, IC<sub>50</sub><br>(ng/mL) |  |  | Site<br>on<br>GP | GP<br>mutations | IgG therapy<br>in mice<br>(EBOV), %<br>survival | Antibody<br>source |
| --- | --- | --- | --- | --- | --- | --- | --- | --- | --- | --- | --- |
|  | EBOV<br>GP | BDBV<br>GP | SUDV<br>GP | EBOV<br>sGP* | EBOV | BDBV | SUDV |  |  |  |  |
| EBOV-293 | 49 | 198 | 91 | 3 | 1,672 | 1,892 | 10,456 | GC | T240A,<br>W275A | 100* | This study |
| EBOV-296 | 39 | 78 | 514 | 4 | 5,758 | 534 | 39,217 | GC | W275A | 20* | This study |
| BDBV-329 | > | 135 | > | 21<br>(BDBV<br>sGP) | > | 770 | > | GC | ND | ND | Flyak *et al.*<br>2016 |
| BDBV-289 | 29 | 20 | 103 | 1 | 602 | 32 | > | GC | W275A,<br>Y241A | 80 | Flyak *et al.*<br>2016 |
| BDBV-43 | 22 | 29 | 21 | 18<br>(BDBV<br>sGP) | 7,248 | 139 | 318 | GC | L273P | 20 | Flyak *et al.*<br>2016 |
| EBOV-237 | 25 | 3,367 | > | 162 | 780 | ND | ND | GC | I260R,<br>N278A,<br>P279A,<br>S322G | 80-90** | Williamson<br>*et al.* 2019 |
| EBOV-437 | 3 | 11 | 4 | 1 | 8,660*** | > | > | GC | W275A | ND | Gilchuk *et al.* 2018 |
| EBOV-442 | 1 | 3 | 6 | 1 | 467 | 1,489 | <75%*** | GC | W275A,<br>L273P | 80 | Gilchuk *et al.* 2018 |
| EBOV-548 | 6 | 7 | 132 | 1 | 1,601 | 2,262 | <12%*** | GC | T240A,<br>R266A,<br>T269A,<br>T270A,<br>I274A,<br>W275 | 40 | Gilchuk *et al.* 2020 |
| 13C6 | 20 | > | > | 2 | > | > | > | GC | T270A,<br>K272A,<br>T240N | 100 | Wilson *et al.* 2000 |
| 2G4 | 203 | > | > | > | 139 | > | > | Base | C511A,<br>G553A,<br>N550A,<br>D552A,<br>C556A,<br>Q508R | 60 | Qiu *et al.*<br>2012 |

We then analyzed all ten glycan cap mAbs for binding enhancement of the base-region-directed mAbs EBOV-515 or EBOV-520 using an approach described previously (Gilchuk et al., 2020). Synergy for each glycan cap antibody followed similar patterns for EBOV-515 and EBOV-520, although enhancement of EBOV-520 binding appears higher likely due to differences in the molecular nature of the epitope (**Fig. 1A**). A steady range of synergistic patterns from no enhancement (for 13C6) to binding nearly equivalent to cleaved GP (GP_CL_) (for EBOV-237) were observed (**Fig. 1A**). It should be noted that BDBV-329 and EBOV-237 are monospecific for the autologous virus BDBV or EBOV, respectively (**Table 1**).

**Figure 1.**
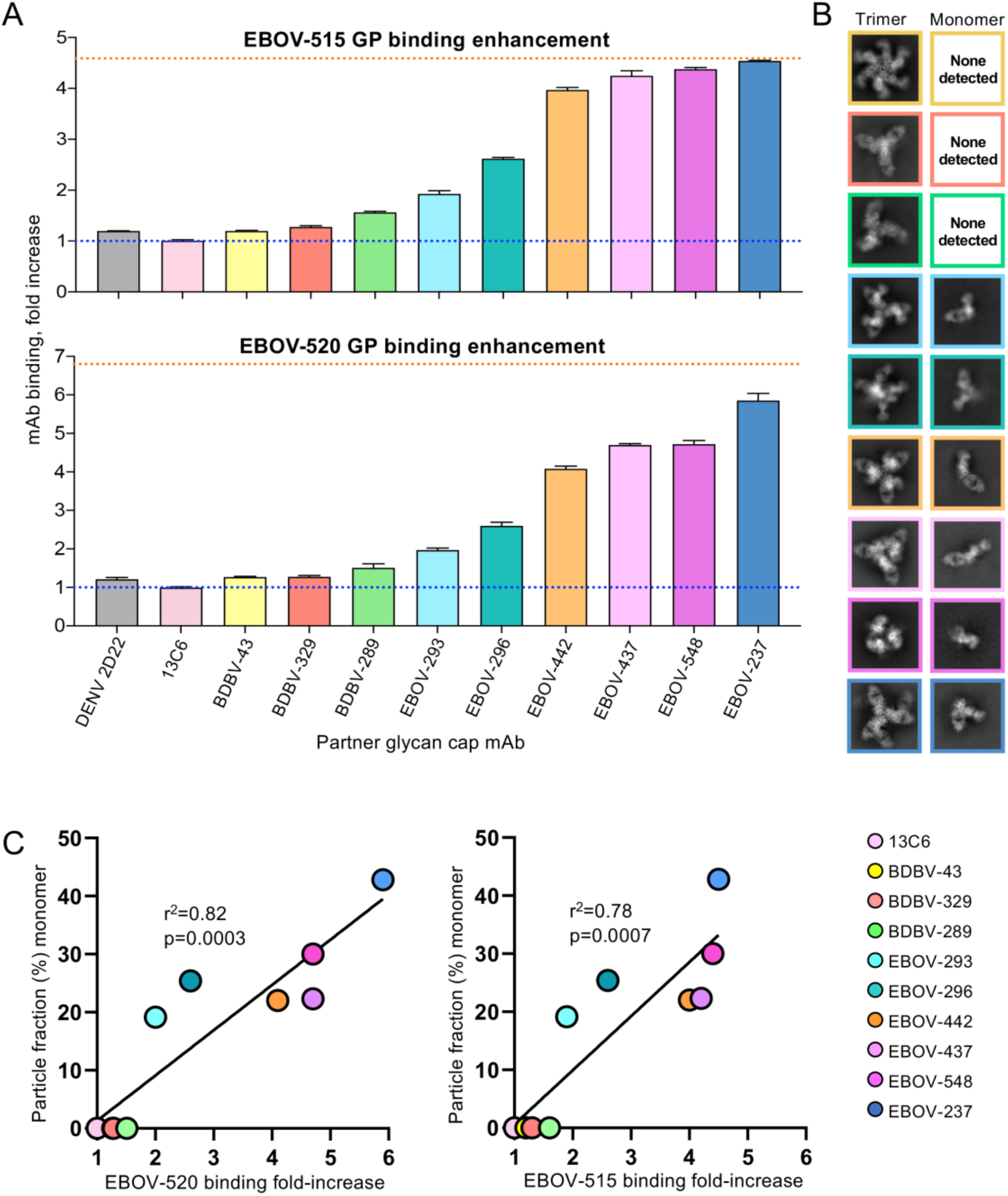
Glycan cap antibody synergy and GP destabilization. **A)** Jurkat cell-surface displayed EBOV GP binding was assessed using fluorescently labeled EBOV-515 or EBOV-520 after prior incubation of cells with individual unlabeled glycan cap antibodies. The blue dotted line represents basal binding of base antibodies without glycan cap antibodies. The orange dotted line represents maximal binding of base antibodies to cleaved GP. Data shown as mean ± SD of technical triplicates **B)** Negative stain 2D-class averages of GP complexes bound to glycan cap antibodies and EBOV-515 demonstrating examples of intact, trimeric complexes (left) and monomeric complexes (right). **C)** Correlation analysis of antibody synergy and GP destabilization by glycan cap antibodies.

We noticed in several of our 2D classes of glycan cap antibody complexes that the GP trimer fell apart into GP monomers, similar to what we had observed with our previous characterization of EBOV-548 (Gilchuk et al., 2020). The amount of GP trimer destabilization was variable across all the complexes, with some antibodies inducing a large amount of GP monomers and others only a stable GP trimer. We specifically avoid inclusion of monomers during protein purification to obtain a pure fraction of trimeric GP as starting material. We therefore hypothesized that glycan cap antibodies destabilize trimers, which in turn may contribute to their synergistic ability with base-region-binding antibodies.

To quantify GP destabilization, we analyzed cryo-EM data collected on glycan cap antibody complexes for structural analysis (*see below*). We also included previous data collected on EBOV-548 complexed with GP (Gilchuk et al., 2020). Particles were selected using a difference of gaussian approach which would not discriminate trimeric complexes from monomeric ones and then performed reference-free 2D classification (**Fig. 1B**). All trimeric and monomeric particles were subsequently subclassified and particle counts were used to determine the percentage of monomeric particles in the cleaned stack of total particles for each dataset.

When plotting the proportion of monomers formed in the presence of each glycan cap antibody, we noted that the amount of destabilization correlated with the extent of antibody synergy (**Fig. 1C**). Antibodies that did not synergize with base-region-binding antibodies displayed little to no destabilization, such as 13C6, BDBV-43, BDBV-329 and BDBV-289. As synergy increased, we saw increasing amounts of destabilized trimers, with EBOV-237 demonstrating the highest level of destabilization (**Fig. 1C**).

### Conservation of a structural β-hairpin motif across synergistic glycan cap antibodies

To determine the structural basis of neutralization and synergy behind glycan cap antibody-based enhancement of base-region-binding antibodies, we solved eight structures of glycan cap antibodies in complex with mucin deleted (ΔMuc (**Fig. 2A-B**, **Fig. S2, Table S2**). Antibodies exhibited a range of angles-of-approach to GP, from obtuse, such as EBOV-437, to nearly parallel to the viral surface, like EBOV-237 (**Fig. 2C**). Additionally, the antibodies are spaced across the surface of GP inversely related to their angle-of-approach (**Fig. 2C**).

**Figure 2.**
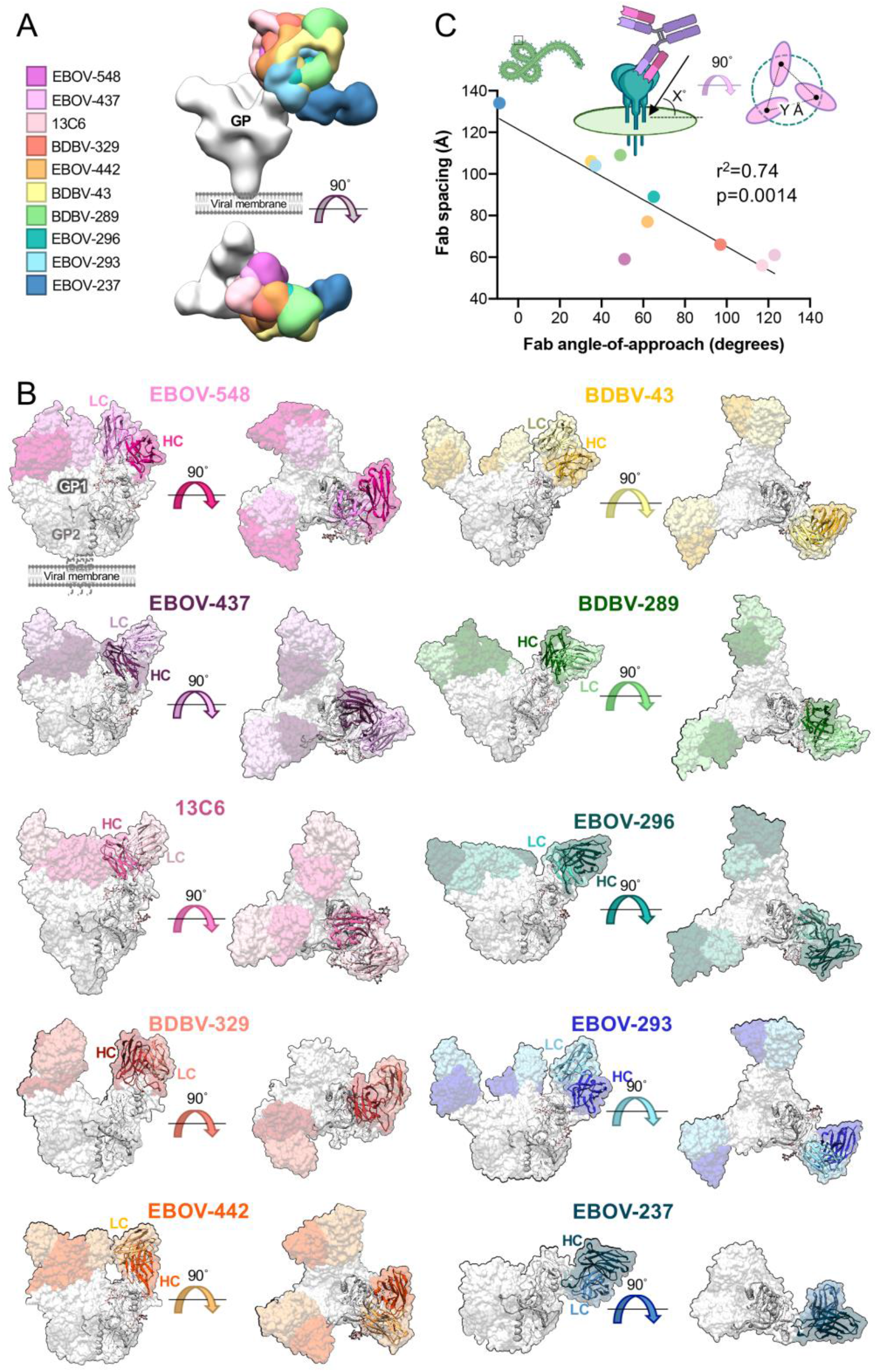
Neutralizing and synergistic glycan cap antibodies bind GP across a wide range of orientations. **A)** Low-pass filtered glycan cap Fabs from cryo-EM structures solved in this study, as well as elsewhere, bound to EBOV GPΔMuc are overlaid to compare binding epitope and angle-of-approach. **B)** Surface representations of cryo-EM structures solved in this study with a fitted ribbon model protomer. Shown are side (left) or top (right) views with respect to the viral membrane. Fab HC is colored in dark tones and LC in light tones. Co-binding antibodies were removed from reconstructions for clarity. **C)** Relationship between antibody angle-of-approach and Fab spacing. An angle-of-approach of zero degrees is considered parallel and 90° is considered perpendicular to the viral surface. An angle-of-approach greater than 90° indicates antibodies that bind inward toward the head domain while less than 0° indicates antibodies that bind upward from the viral membrane. Fab spacing is determined by averaging the distance from the same point on modeled Fab hinge terminal residues in the HC and LC. Antibodies are labeled as in part A. See also Figure S2 and Figure S4.

Resolutions achieved for glycan cap antibody cryo-EM structures ranged from 3.3 to 4.4 Å for six of our complexes (**Table S2, Fig. S2A-F**); however, preferred orientation, sub-stoichiometric Fab binding and trimer instability resulted in limited resolution for BDBV-329 and EBOV-237 bound structures (**Fig. S2G-H**). We did, however, model BDBV-329 where resolutions ranged from 4 to 5 Å at the antibody binding interface (**Fig. S2G**). The local resolutions for the EBOV-237 structure were particularly poor, and we therefore chose only to dock a homology model for interpretation (**Fig. S2H**). Most of our structures were determined in complex with EBOV-515 in order to assist with angular sampling and alignment, but we chose not to model EBOV-515 and removed this density from our figures for clarity and to focus on glycan cap antibodies (**Fig. 3A-B**).

**Figure 3.**
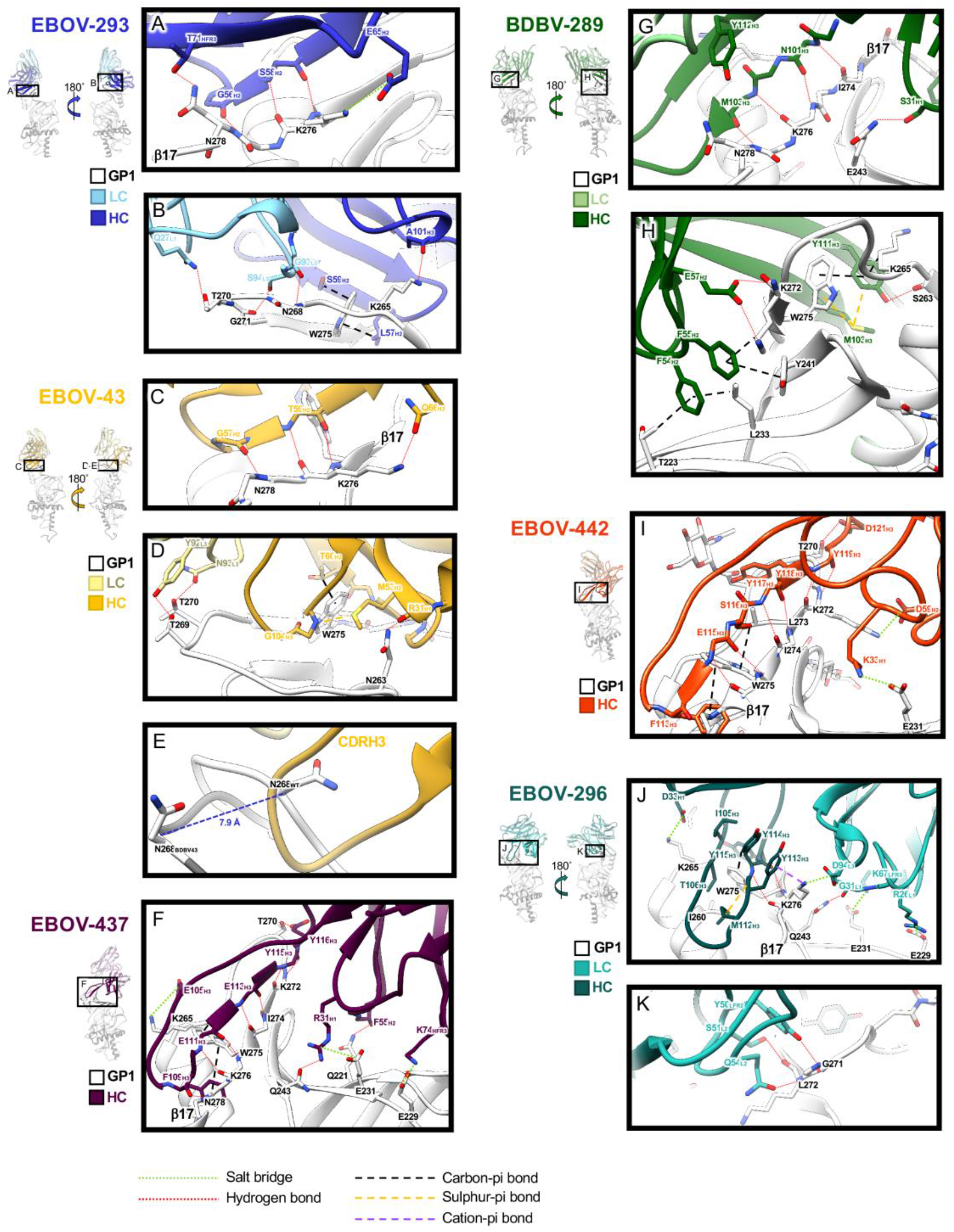
Structural details of glycan cap antibody binding to GP. Single protomers from structural models are shown with close-up views of interacting regions. HCs are rendered in darker colors and LCs in lighter colors, with GP1 colored white. Important residues that coordinate interaction and binding are highlighted. **A)** Key residues in the EBOV-293 CDRH2 hydrogen bond along the length of β17 with an additional potential salt bridge between E65_H2_ and K276_GP1_. **B)** The EBOV-293 CDRH2 and H3 make additional contacts, including at W275, and the LC forms potential hydrogen bonds between α2 and β17. **C)** Similar to EBOV-293, the BDBV-43 CDRH2 loop binds along β17. **D)** The BDBV-43 CDRH2 makes additional contacts at W275 and also contacts the loop between α2 and β17 via its LC. **E)** The EBOV-43 CDRH3 loop displaces the loop between α2 and β17, shifting N268 by ~8 Å (apo-GP in white and BDBV-43 bound GP in grey). **F)** EBOV-437 makes contact with GP exclusively with its HC, hydrogen bonding along β17 with its CDRH3 and contacting the head domain in several places. **G)** BDBV-289 makes extensive hydrogen bonds with its CDRH3 along β17. **H)** The BDBV-289 CDRH3 contacts W275 via methionine-aromatic and pi-pi interactions. Additional contacts are made with the head domain of GP via hydrophobic interactions with the CDRH2. **I)** BDBV-442 makes contact with GP exclusively with its HC. The CDRH3 makes hydrogen bonds along β17, with W275 with hydrophobic interactions and along the loop between α2 and β17. **J)** EBOV-296 binds to GP along β17, contacting W275 via methionine-aromatic and pi-pi interactions. The LC makes further contact to the head domain of GP with several potential salt bridges. **K)** The EBOV-296 LC also makes contact to the loop between α2 and β17. See also Figure S3 and Figure S4, Table S2 and Table S3.

All glycan cap antibodies make contacts exclusively within GP1 and are heavily biased toward HC contacts (**Fig. 3**, **Table S3, Fig. S3**). Antibody contacts are focused on the β17 strand of GP1 from residues 268-280 with a majority of contacts centered around W275 (**Fig. 3**, **Table S3**), which when mutated to alanine abrogates binding (**Table 1, Fig. S1B**). Additionally, most glycan cap antibodies make some contact with the inner head domain (**Fig. 3**, **Table S3, Fig. S3**). These contacts are characterized by hydrogen bonding along the length of β17 with either short CDRH2 loops for EBOV-293 (**Fig. 3A-B**, **Fig. S3A**) and BDBV-43 (**Fig. 3C-D**, **Fig. S3B**) or extended, long (≥21 amino acids) CDRH3 loops for EBOV-437 (**Fig. 4F**, **Fig. S5C**), BDBV-289 (**Fig. 3G-H**, **Fig. S3D**), EBOV-442 (**Fig. 3I**, **Fig. S3E**) and EBOV-296 (**Fig. 3J-K**, **Fig. S3F**), very similar to EBOV-548 as we previously reported (Gilchuk et al., 2020). Outside of the hydrogen bonding that occurs along β17, several glycan cap antibodies make additional stabilizing bonds, including hydrogen bonds, salt bridges, carbon-pi and pi-pi bonds with other portions of GP1 (**Fig. 3**). Methionine-aromatic interactions also appear in several of the glycan cap antibodies, particularly with W275 in GP1 (**Fig. 3D, H, J**). These types of interactions are thought to provide additional stability compared to purely hydrophobic interactions, can act at long distances (~5-6 Å) and are thought to be less sensitive to changes in the local environment (Valley et al., 2012), which may help contribute to increased breadth.

**Figure 4.**
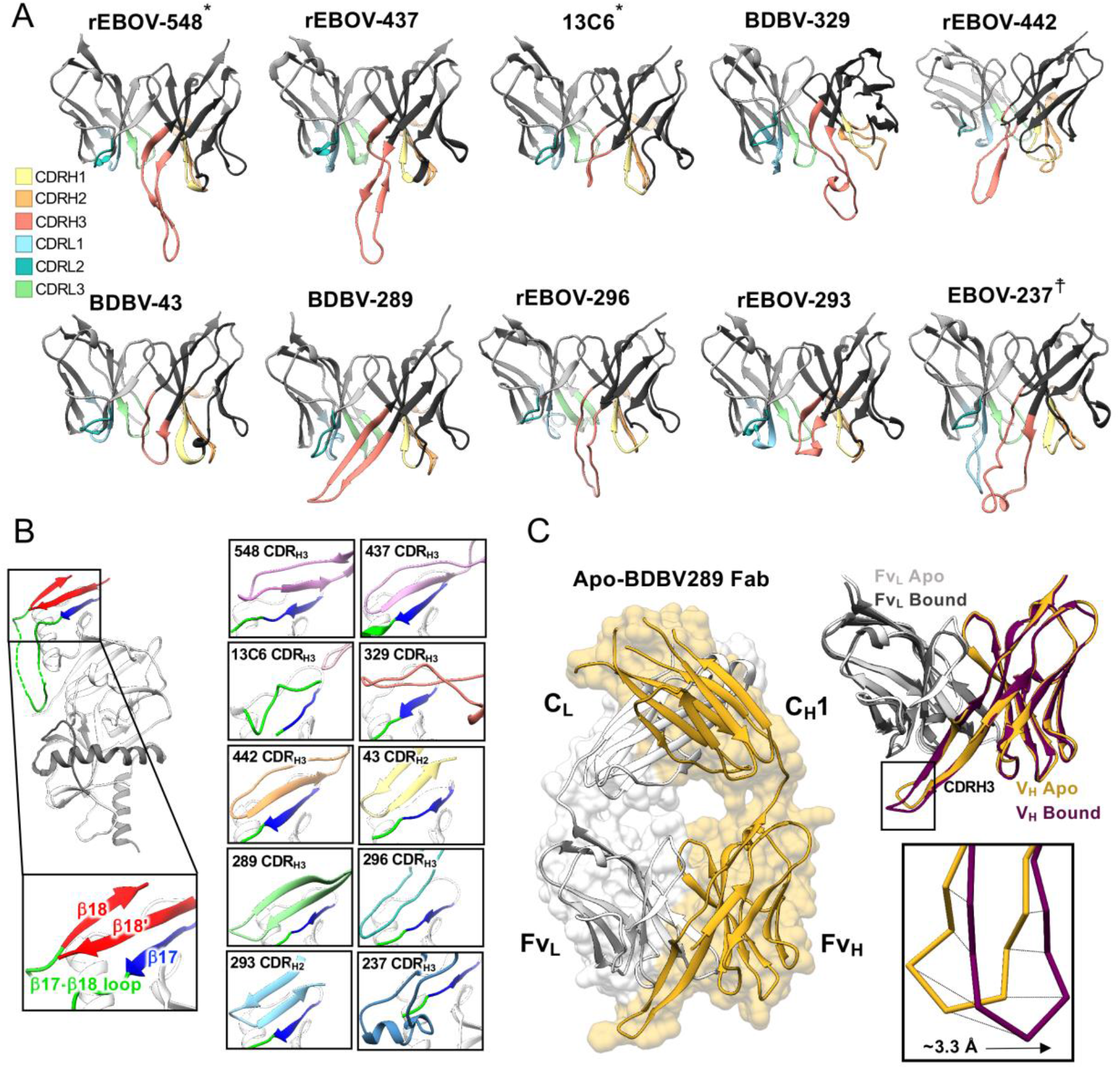
Glycan cap antibody paratopes feature long CDRH3 or short CDRH2 loops with beta-hairpin structures that mimic and displace the β18-β18′ region in the glycan cap. **A)** Ribbon models of the glycan cap antibody Fv domains with CDR loops highlighted. The HC is in dark gray (right) and the LC is in light grey (left). **B)** Structures highlighting the interaction of each of the glycan cap antibodies with the β17 strand, which forms the basis of an extended beta sheet in the glycan cap with the β17-β18 loop and β18-β18′ hairpin motif (shown on the left). **C)** Crystal structure of BDBV-289 Fab. Shown on the right is comparison of the apo- and GP-bound forms of BDBV-289. *From a previous study; ☨shown as an initial homology model. EBOV-548 (PDB 6UYE) and 13C6 (PDB 5KEL) are included in this figure for comparison. See also Figure S5, Table S2 and Table S4.

The CDRH3 loops of the glycan cap antibodies generally adopt an extended β-hairpin motif with either partial or full β-strand secondary structure (**Fig. 4A**). These loops also pair with β17 in GP1 to form an extended β-sheet and displace the β18-β18′ hairpin by mimicking its structure, as was observed in our previous structure of EBOV-548 (**Fig. 4B**). BDBV-43 and EBOV-293 alternatively use shorter CDRH2 loops to pair with β17 (**Fig. 4A-B**). Conversely, 13C6 has a much shorter CDRH3 loop and does not make full contact with β17 (**Fig. 4B**), possibly explaining its lack of synergy with base antibodies (**Fig. 1A**). EBOV-237 and BDBV-329 are unique among the antibodies we examined here owing to very long CDRH3 loops at 25 or 33 amino acids, respectively.

We also determined the unliganded crystal structure of BDBV-289 Fab to 3.0 Å resolution to compare the confirmations of the CDR loops prior to GP engagement (**Fig. 4C**, **Table S4**). The structure of the unliganded BDBV-289 Fab is very similar to BDBV-289 Fab bound to EBOV GPΔMuc, with an RMSD of 1.6 Å for the Fv portions of the HC and LC (**Fig. 4C**). There is a slight shift of the CDRL3 to accommodate the α2-β17 loop in the glycan cap, and a larger shift of CDRH3 (**Fig. 4C**). In the GP-bound structure, the CDRH3 loop moves toward GP by an average distance of ~3.3 Å (**Fig. 4C**). In the crystal structure, this movement is blocked by a crystal lattice interaction, but this difference may indicate flexibility in the tip of this loop.

### The β18-β18′ hairpin anchors the mucin-like domains and shields a hydrophobic patch in the glycan cap

The β18-β18′ region of the glycan cap forms a β-hairpin that anchors the MLD, forming an extended beta sheet with the underlying core of GP1 (**Fig. 5A**) (Zhao et al., 2016). Due to the recurrence of the β18-β18′ hairpin epitope within the glycan cap and its role in anchoring down MLD, we have named this portion of the glycan cap the “MLD-anchor” (**Fig. 5A**). Upon binding of glycan cap antibody, the MLD-anchor is displaced, revealing a patch of hydrophobic residues, which we refer to as the “MLD-cradle” (**Fig. 5B**).

**Figure 5.**
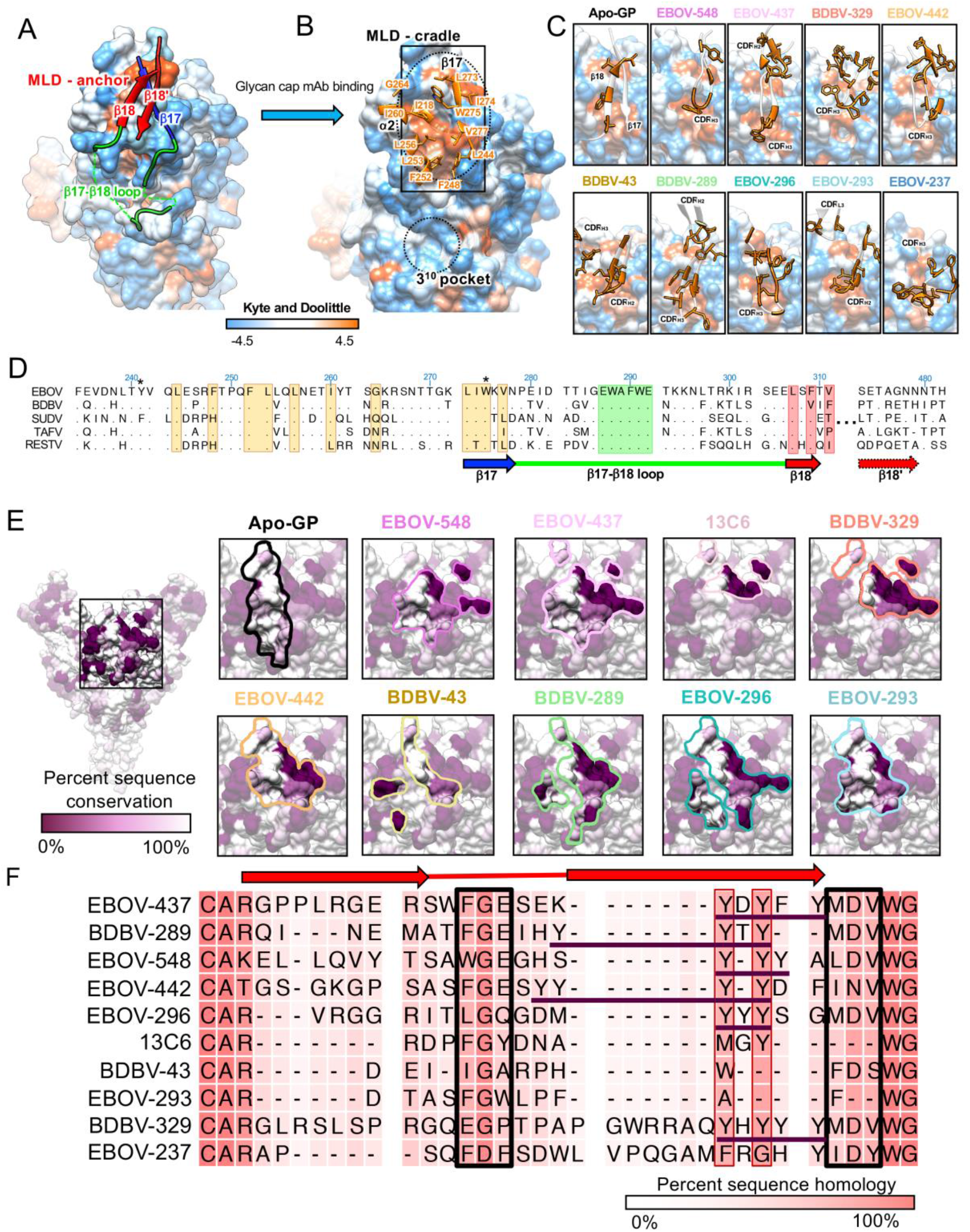
Glycan cap antibodies target a conserved, hydrophobic cradle that anchors the mucin-like domains to GP1. **A)** Hydrophobicity surface rendering of apo-EBOV GP protomer (PDB 5JQ3) with the MLD-anchor (β18-β18′) highlighted in red. Using the Kyte and Doolittle scale (Kyte and Doolittle, 1982), hydrophobic residues are colored orange with hydrophilic ones in blue. **B)** Upon glycan cap mAb binding, the MLD-anchor is displaced, exposing a hydrophobic pocket we term the “MLD-cradle”. The cradle lies within a groove formed by α2 and β17, directly above the 3^10^ pocket. Key residues of the cradle are indicated. The MLD-cradle is composed of residues from α2 and β17 as well as some additional residues that lie deeper in the core of GP1, including I218, F248, F252, L253, L256, I260, G264, L273, I274, W275, V277 and L244. The cradle is segmented in the middle by W275, which may explain this residue’s pivotal role in the binding of many glycan cap antibodies to GP. **C)** Interaction of glycan cap mAb HC loops with the MLD-cradle (from the rectangle in panel B). Key hydrophobic residues from antibody paratopes are indicated. **D)** Sequence alignment of the MLD-anchor and cradle epitope for the five main ebolaviruses (EBOV Q05320, BDBV B8XCN0, SUDV Q66814, TAFV Q66810 and RESTV Q66799) with topology indicated below. Residues highlighted in orange are key hydrophobic residues that form the cradle, those in green form the base of the β17-β18 loop that interact with the base of the fusion loop, and those in red are key residues from β18 that interact with the cradle in apo-GP. Those marked with a * are common escape mutants for this epitope. **E)** Glycan cap antibody footprints highlighted on the structure of apo-GP colored to reflect conservation, with dark purple indicating complete lack of conservation and white indicating complete conservation. **F)** Shown is a sequence alignment of the CDRH3 region from each of the glycan cap antibodies analyzed in this study, with darker pink indicating complete conservation and light pink indicating complete lack of conservation. The beta-turn-beta structure common to these paratopes is indicated above. Key sequences that are similar among these antibodies are boxed in black with “Y” stretches from *IGHJ6* gene usage underlined in purple. See also Figure S5.

The MLD-anchor contains complementary hydrophobic residues along the β18′ strand that are buried by the MLD-cradle (**Fig. 5C**). Through molecular mimicry, the CDRH3 or CDRH2 loops of each of the neutralizing glycan cap antibodies characterized in this study bury analogous hydrophobic residues in the cradle, displacing the anchor (**Fig. 5C**). Our structures of EBOV-548 (Gilchuk et al., 2020) Fab and BDBV-289 Fab bound to GP indicate that although binding abrogates attachment of the MLD-anchor, the β17-β18 loop most likely remains tethered to the base of the IFL via W291_GP1_ to N512_GP2_. However, glycan cap binding may remove some restraint on the β17-β18 loop, allowing increased binding by GP base-directed antibodies.

The sequence of the N-terminal portion of the MLD-anchor (β18), the MLD-cradle and the β17-β18 loops are relatively conserved throughout all ebolaviruses, however, the surrounding regions in the glycan cap are not (**Fig. 5D-E**, **Fig. S4**). The glycan cap antibody contacts described here are focused toward β17 but other contacts outside this region are also observed (**Fig. S4, Table S3**). While each glycan cap antibody makes contacts outside the most conserved regions, there is less reliance on these regions for contact in the more cross-reactive antibodies (**Fig. S4**).

### Germline analysis and conservation of features within glycan cap antibody paratopes

The glycan cap antibodies described here share several common features, including a majority (5 out of 9) deriving from the *IGHV1-69* germline gene segment (**Table S1**). Frequent use of the *IGHV1-69* gene is common in the antibody repertoires of those infected by influenza virus (Lang et al., 2017), HCV (Chan et al., 2001), HIV-1 (Huang et al., 2004), and other pathogens (Chen et al., 2019). The *IGHV1-69* gene is thought to be superior for viral neutralization at certain epitopes due to the presence of key germline encoded hydrophobic residues, especially in the CDRH2, as well as for breadth due to a large repertoire of allelic and copy number variations (Chen et al., 2019). Despite a wide range of donors, we found these characteristics present in the ebolavirus antibodies described here (**Table S1, Fig. S5**).

Eight of the nine of the human antibodies described here use *IGHV1-69* and/or *IGHJ6* genes to form their HCs (**Table S1, Fig. S5**), the exception being EBOV-237. The use of *IGHV1-69* imparts a germline-encoded CDRH2 with several hydrophobic residues, which is used by BDBV-43 and EBOV-293 to bind to the MLD-cradle (**Fig. 5C**). In these cases, the CDRH3 loops are shorter (**Fig. S5**). For the rest of the glycan cap antibodies, usage of *IGHJ6* appears to be key, due to the presence of a patch of tyrosine residues in the germline gene which sit on the C-terminal end of the CDRH3 (**Fig. S5**). Overall, somatic hypermutation (SHM) was generally high throughout all glycan cap mAbs studied here, with an average of ~11% or ~6% amino change from germline for the V_H_ or V_L_ regions, respectively (**Table S1**).

Despite varying CDRH3 length, the tip of the CDRH3 hairpin contains a highly conserved glycine surrounded by hydrophobic residues and a C-terminal tyrosine motif (**Fig. 5F**). This glycine and hydrophobic tip help to insert the CDRH3 loop into the MLD-cradle (**Fig. 5A-C**) and assists in formation of the hairpin structure necessary for proper binding. The C-terminal tyrosine motif stabilizes longer CDRH3 loops within the core of the paratope and provides additional, non-specific hydrophobic contacts within the core of the epitope.

### Glycan cap antibodies inhibit cleavage

The underlying molecular mechanism for how an antibody neutralizes is related to its ability to inhibit viral infection, which can be achieved by diverse mechanisms including cleavage inhibition. To determine the ability of the antibodies used in this study to inhibit cleavage, we performed a cleavage-blocking assay as previously described (Gilchuk et al., 2018) (**Fig. 6A**). Jurkat cells stably transduced with EBOV GP (Jurkat-EBOV GP) were pre-incubated with individual antibodies followed by treatment with thermolysin to mimic cathepsin cleavage to yield membrane-displayed GP_CL_ (Jurkat-EBOV GP_CL_). The exposure of the receptor binding site (RBS) on GP_CL_ was measured by the level of binding of fluorescently labeled RBS-specific mAb MR78 that does not bind uncleaved EBOV GP (Flyak et al., 2015) The epitope of glycan cap antibodies is being removed by cleavage, and in a separate assay we confirmed that none of tested antibodies, except EBOV-442, compete with MR78 on Jurkat-EBOV GP_CL_ (**Fig. S6**). EBOV-442 partially competed with MR78 (**Fig. S6**), suggesting incomplete removal of its epitope by thermolysin that may have a minor effect on quantification of cleavage inhibition by this antibody. All EBOV GP-reactive glycan cap antibodies revealed dose-dependent cleavage inhibition and most of them fully blocked cleavage at the highest tested concentration of 60 μg/mL (**Fig. 6A**). Base antibody 2G4 and control antibody 2D22 did not inhibit cleavage. Although the glycan cap antibodies in this study do not interact directly with the cathepsin cleavage loop, the disruption or dislocation of the MLD may provide an obstacle for the recognition or cleavage activity by enzymes (**Fig. 6B**).

**Figure 6.**
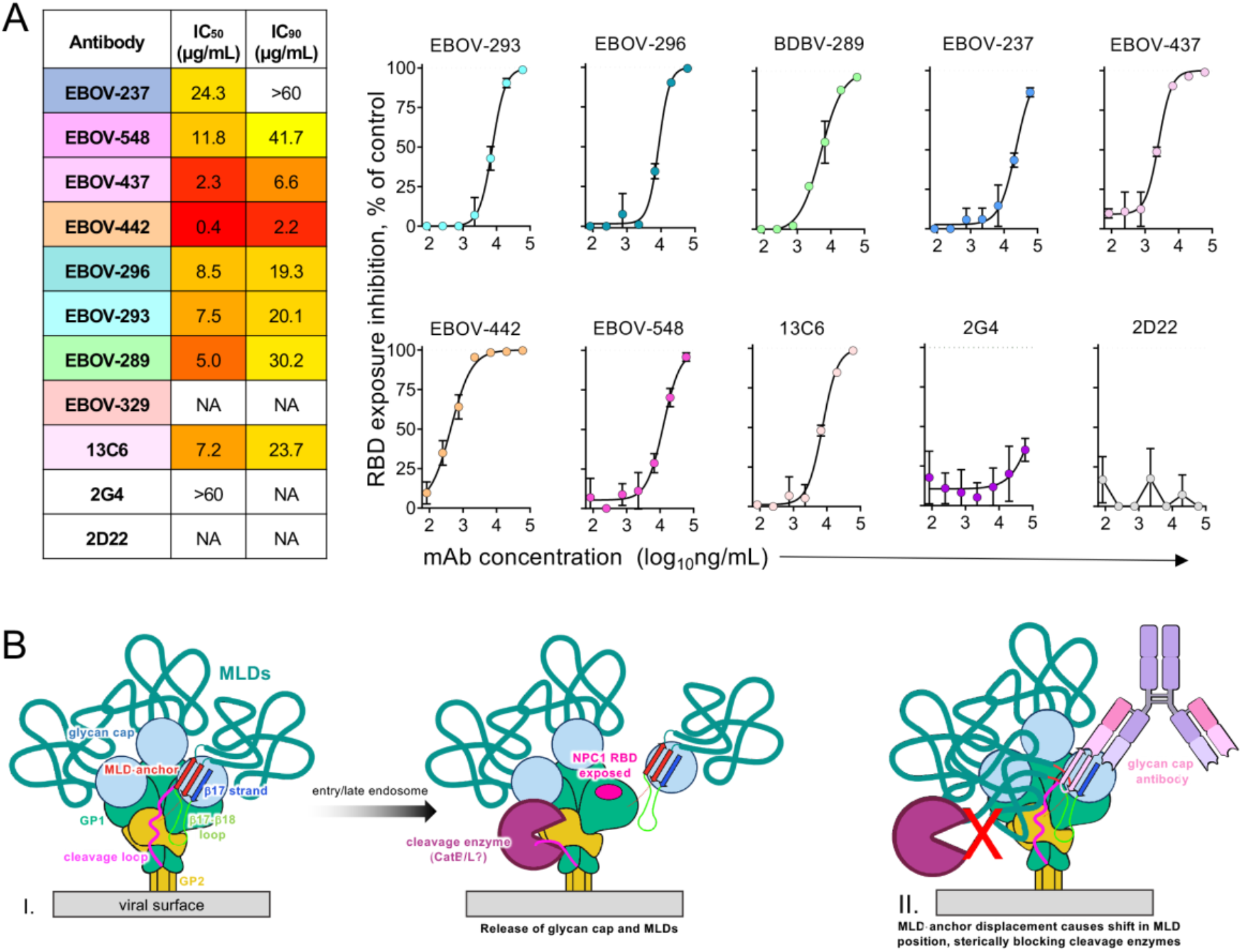
Cleavage inhibition by glycan cap antibodies. **A)** Jurkat-EBOV GP was incubated with various concentrations of antibodies, treated with thermolysin, and then assayed using flow cytometry for exposure of the receptor binding site (RBS) as measured by binding of a fluorescently labeled MR78 antibody that recognizes the RBS. Determined IC_50_ and IC_90_ values (left) and dose-dependent inhibition curves (right) are shown. Dotted line indicates % RBS exposure in the presence of 2D22 control. BDBV-329 was excluded because it does not bind to EBOV GP, and BDBV-43 was not tested due to poor recombinant expression. **B)** Proposed model of the GP inhibition by glycan cap antibodies: I. Enzyme cleavage of a loop draped over the outside of GP (magenta) is thought to release the glycan cap and attached MLD. II. Glycan cap antibodies that bind to the MLD-cradle displace the MLD-anchor and thus the MLDs themselves, potentially shifting their position and sterically blocking access to the cleavage loop by enzymes, especially on a GP-dense viral surface. See also Figure S6.

## Discussion

Our previous structure of the EBOV-548/EBOV GP complex first revealed the glycan cap binding site containing the β18-β18′ hairpin (Gilchuk et al., 2020); however, the extensive structural evidence we provide here more completely describes this epitope, which we coin the MLD-anchor and cradle. The displacement of the MLD-anchor suggests that it is bound transiently, similar to the β17-β18 loop (West et al., 2019). Anecdotally, we and others have often noticed that the glycan cap is not well resolved in negative stain and cryo-EM structures of GP that lack coordinating glycan cap antibodies, suggesting this entire domain may be loosely attached to GP. This transient structural feature may aid in removal of the glycan cap upon cleavage for exposure of the NPC1 binding site. The MLD-anchor makes very limited contact with the underlying hydrophobic cradle, essentially mediated by a single β-strand. These characteristics have been observed for other antibodies that bind with hydrophobic, hairpin CDR loops, suggesting a conserved mechanism for neutralization that extends to other viruses (Lee et al., 2017; Pancera et al., 2010; Yuan et al., 2019).

Some of the antibodies in this study shared structural features that correlated with shared antibody germline gene usage, despite having been isolated from separate donors. For example, BDBV-43 and EBOV-293 are both encoded by the *IGHV1-69*\**09* gene allele, which is known to contain a germline-encoded hydrophobic CDRH2. These antibodies, accordingly, use their CDRH2 loops to access β17. However, several other antibodies in this study also use the *IGHV1-69* gene, but alternatively use longer CDRH3 loops to access β17. BDBV-43 and EBOV-293 also share almost identical LC usage (*IGKV3-15*\**01* and *3-20*\**01* for V_L_ genes and *IGKJ5*\**01* for J_L_ genes, respectively). Their structures are nearly identical as well, which indicates that LC pairing may influence access to β17 by *IGHV1-69* gene-encoded CDRH2 loops. However, this barrier can be overcome through a larger number of non-templated (N) nucleotide additions within the CDRH3 when using alternative LC pairing. These observations may indicate that germline targeting vaccines could be an effective strategy for eliciting similar antibodies for EBOV in healthy individuals, similar to what has been shown in HIV-1 vaccine studies (Steichen et al., 2016; Steichen et al., 2019).

The glycan cap antibodies described here generally have high levels of SHM, with EBOV-293 containing 24 mutations from the inferred germline gene in its heavy chain (**Table S1**). This count also does not consider potential somatic mutations in the long CDRH3 loops, whose germline origins cannot be predicted but likely arise from large numbers of N-additions during the original V-D-J recombination event. How glycan cap antibody SHM compares to mutation frequency in antibodies directed toward other epitopes is not well explored, but the amount of SHM we observe for these neutralizing glycan cap mAbs is higher than is generally reported in EBOV survivor repertoires (Davis et al., 2019). Glycan cap antibodies are now known to form a large portion of the adaptive response to natural infection (Bornholdt et al., 2016b; Flyak et al., 2016; Wec et al., 2017). Several of these antibodies can potently neutralize, however, they are often mono-specific. It is unclear how the smaller subset of rarer, broadly neutralizing glycan cap antibodies develops. Our observations indicate that they may require higher levels of SHM combined with structural adaptations in order to reach cryptic epitopes shielded by the MLD, the MLD-anchor and glycans.

The mechanism of viral neutralization by glycan cap antibodies is unclear. Potentially, these antibodies could act indirectly by preventing access to a cleavage loop that is necessary to cleave during viral entry (Bornholdt et al., 2016a) (**Fig. 6B**, **part I**). The MLDs are large, accounting for over half of the mass of GP, unstructured and highly glycosylated. While the MLDs on ebolaviruses are known to sit atop the GP, those on marburgviruses are thought to drape over the sides (Hashiguchi et al., 2015). This difference may occur because marburgviruses lack the structured glycan cap that is found in ebolaviruses (King et al., 2018). Consequently, the NPC1 receptor binding site is fully exposed on full-length GP in marburgviruses (Flyak et al., 2015), while it is hidden under the glycan cap and MLD on ebolaviruses. Therefore, the MLD-anchor appears to pin the MLD down to the top of ebolavirus GPs, keeping them above the GP and out of the way of the cleavage loop. Displacing the MLD-anchor may displace the MLDs themselves while retaining covalent attachment of these large domains to GP (**Fig. 6B**, **part II**). Within the dense environment of the ebolavirus surface, in which many GP spikes are known to crowd together in close proximity (Tran et al., 2014), this displacement may cause the MLD to drape over the cathepsin cleavage loops, thus blocking access by enzymes.

Overall, our data collectively provide the molecular basis for breadth of reactivity and virus neutralization by potent glycan cap antibodies and suggest a rational strategy for the design of broad therapeutic antibody cocktails.

## Acknowledgements

This work was supported by NIH grants U19 AI109762 and U19 AI142785. T.A. is supported by a Kellogg Graduate Student Fellowship from Scripps Research. We would like to thank the Joint Center for Structural Genomics at Scripps Research and Henry Tien for assistance with setting up crystal trays. We like to thank Dr. Robyn Stanfield for assistance with looping and shipping crystals, for assistance with collecting X-ray diffraction data, reducing the data and phasing the crystal structure. We also thank Dr. Ian Wilson for generously sharing synchrotron time to collect X-ray diffraction data. Use of the Stanford Synchrotron Radiation Lightsource, SLAC National Accelerator Laboratory, is supported by the U.S. Department of Energy, Office of Science, Office of Basic Energy Sciences under Contract No. DE-AC02-76SF00515. The SSRL Structural Molecular Biology Program is supported by the DOE Office of Biological and Environmental Research, and by the National Institutes of Health, National Institute of General Medical Sciences (including P41GM103393). The contents of this publication are solely the responsibility of the authors and do not necessarily represent the official views of NIGMS or NIH.

The Jurkat-EBOV GP cell line was a kind gift form Carl Davis and Rafi Ahmed. We thank Hannah Turner, Bill Anderson, Jonathan Torres, Gabriel Ozorowski and Charles Bowman from Scripps Research for their assistance with cryo-EM sample prep, microscope operation, data collection and processing.

## Author contributions

C.D.M. performed protein production for all cryo-EM and kinetic experiments, performed cryo-EM experiments and analysis, crystallized the BDBV289 Fab and performed kinetic experiments and analysis. C.D.M. collected the crystal data and phased the data. J.F.B. built and validated the crystal structure. J.C. produced antibody Fab for the crystallography trials. P.G. performed synergy, cleavage assays and assembled information for Table 1. T.A. helped to perform GP stability assays and collect cryo-EM data. L.W. expressed and purified EBOV-237 and BDBV-329 Fab. P.A.I., K.H., N.K., X.S., A.I.F. and A.B. isolated and characterized EBOV-293 and EBOV-296. A.L.B., E.D. and B.J.D. performed alanine scanning and characterization of EBOV-293 and EBOV-296. C.D.M., P.G., A.B., J.E.C. and A.B.W. designed the experiments. C.D.M. wrote the manuscript.

## STAR Methods

### RESOURCE AVAILABILITY

#### Lead contact

Further information regarding requests for resources and reagents should be directed to and will be fulfilled by the Lead Contact, Andrew Ward.

#### Materials availability

Plasmids generated in this study are available upon request by the Lead Contact.

#### Data and code availability

The cryo-EM maps and structural coordinates generated during this study are available at the Electron Microscopy Data Bank (www.ebi.ac.uk/pdbe/emdb) and the Worldwide Protein Data Bank (www.pdb.org). The accession codes for the following cryo-EM maps reported in this paper are: EMD-22839 (EBOV GPΔMuc:BDBV289 Fab), EMD-22841 (BDBV GPΔMuc:BDBV43 Fab and ADI-15878 Fab), EMD-22853 (EBOV GPΔMuc:EBOV-437 Fab and EBOV-515 Fab), EMD-22848 (EBOV GPΔMuc:EBOV-442 Fab and EBOV-515 Fab), EMD-22842 (EBOV GPΔMuc:EBOV-293 Fab and EBOV-515 Fab), EMD-22847 (EBOV GPΔMuc:EBOV-296 Fab and EBOV-515 Fab), EMD-22851 (EBOV GPΔMuc:BDBV-329 Fab and EBOV-515 Fab) and EMD-22852 (EBOV GPΔMuc:EBOV-237 Fab and EBOV-515 Fab). The accession codes for PDB files are: 7KEJ (EBOV GPΔMuc:BDBV-289 Fab), 7KEW (BDBV GPΔMuc:BDBV-43 Fab), 7KFH (EBOV GPΔMuc:EBOV-437 Fab), 7KFB (EBOV GPΔMuc:EBOV-442 Fab), 7KEX (EBOV GPΔMuc:EBOV-293 Fab), 7KF9 (EBOV GPΔMuc:EBOV-296 Fab), 7KFE (EBOV GPΔMuc:BDBV-329 Fab) and 7KFG (unliganded BDBV289 Fab).

### EXPERIMENTAL MODEL AND SUBJECT DETAILS

#### Human samples

Human PBMCs were obtained from a survivor of the 2014 EVD epidemic who acquired the infection in the Democratic Republic of Congo and was treated in the Nebraska Medical Center in the United States. A male human survivor was age 57 when PBMCs were collected. PBMCs were collected after the illness had resolved, following written informed consent. The studies were approved by the Institutional Review Board of Vanderbilt University Medical Center.

#### Cell lines

Suspension adapted HEK293F cells were obtained from ThermoFisher Scientific and cultured in serum-free FreeStyle medium. Cells were maintained in shaking incubators at 100% humidity, 37°C and 8% CO_2_. Expi293F cells (ThermoFisher Scientific) were maintained at 37 °C in 8% CO_2_ in Expi293F Expression Medium (ThermoFisher Scientific). ExpiCHO cells (ThermoFisher Scientific) were maintained at 37°C in 8% CO_2_ in ExpiCHO Expression Medium (ThermoFisher Scientific). The Jurkat-EBOV GP (variant Makona) cell line stably transduced to display respective GP on the surface (Davis et al., 2019) was a kind gift from Carl Davis (Emory University, Atlanta, GA). Jurkat-EBOV GP cells were maintained at 37°C in 8% CO_2_ in RPMI-1640 medium (Gibco) supplemented with 10% fetal heat-inactivated fetal bovine serum (FBS). Mycoplasma testing of Expi293F and ExpiCHO cultures was performed on a monthly basis using a PCR-based mycoplasma detection kit (ATCC). Cell lines were not authenticated following purchase.

#### Viruses

Mouse-adapted EBOV Mayinga (EBOV-MA, GenBank: AF49101) virus was described previously (Bray et al., 1998).

#### Mouse models

Seven-to eight-week old female BALB/c mice were obtained from the Jackson Laboratory. Mice were housed in microisolator cages and provided food and water *ad libitum*. Challenge studies were conducted under maximum containment in an animal biosafety level 4 (ABSL-4) facility of the Galveston National Laboratory, UTMB. The animal protocols for testing of mAbs in mice were approved by the Institutional Animal Care and Use Committee (IACUC) of the University of Texas Medical Branch in compliance with the Animal Welfare Act and other applicable federal statutes and regulations relating to animals and experiments involving animals.

### METHOD DETAILS

#### Isolation of mAbs EBOV-293 and EBOV-296

Hybridoma cell lines secreting human mAbs were generated as described previously (Flyak et al., 2018). In brief, previously cryopreserved samples were transformed with Epstein-Barr virus, CpG and additional supplements. After 7 days, cells from each well of the 384-well culture plates were expanded into four 96-well culture plates using cell culture medium containing irradiated heterologous human PBMCs (recovered from blood unit leukofiltration filters, Nashville Red Cross) and incubated for an additional four days. Plates were screened for EBOV GP antigen-specific antibody-secreting cell lines using enzyme-linked immunosorbent assays (ELISAs). Cells from wells with supernatants reacting with antigen in an ELISA were fused with HMMA2.5 myeloma cells using an established electrofusion technique (Yu et al., 2008). Antibody heavy- and light-chain variable region genes were sequenced from hybridoma lines that had been cloned biologically by single-cell flow cytometric sorting. Briefly, total RNA was extracted using the RNeasy Mini kit (QIAGEN) and reverse-transcriptase PCR (RT-PCR) amplification of the antibody gene cDNAs was performed using the PrimeScript One Step RT-PCR kit (CLONTECH) according to the manufacturer’s protocols with gene-specific primers (Thornburg et al., 2016). The thermal cycling conditions were as follows: 50°C for 30 min, 94°C for 2 min, 40 cycles of (94°C for 30 s, 58°C for 30 s and 72°C for 1 min). PCR products were purified using Agencourt AMPure XP magnetic beads (Beckman Coulter) and sequenced directly using an ABI3700 automated DNA sequencer. The identities of gene segments and mutations from germlines were determined by alignment using the ImMunoGeneTics database (Giudicelli and Lefranc, 2011).

#### Synergistic binding to cell-surface-displayed GP

The assay was performed as described previously (Gilchuk et al., 2018). Briefly, Jurkat-EBOV GP cells were pre-incubated at 4°C for 30 min with individual unlabeled glycan cap-specific mAbs at a saturating for GP binding concentration (20 μg/mL) in PBS containing 2% FBS and 2 mM EDTA, and then Alexa Fluor 647-labeled mAbs EBOV-515 or EBOV-520 were added to a total concentration of labeled mAbs of 10 μg/mL. Cells were incubated at 4°C for additional 2 h, then washed and antibody binding was analyzed by flow cytometry using an iQue Screener Plus flow cytometer (Intellicyt). Controls included binding of labeled mAb to mock-transduced Jurkat cells (background), binding of labeled mAb alone to intact Jurkat-EBOV GP (a baseline level of binding to calculate fold change in a presence of glycan mAb), and binding of labeled mAb alone to cleaved Jurkat-EBOV-GP (maximal saturating binding signal). Results are expressed as fold-increase in median fluorescence intensity (MFI) of labeled mAb binding in the presence of the tested unlabeled mAb minus background signal from mock control.

#### ELISA binding assays

To assess mAb binding at different pH, wells of 96-well microtiter plates were coated with purified, recombinant EBOV, BDBV or SUDV GPΔTM ectodomains or EBOV sGP at 4°C overnight. Plates were blocked with 2% non-fat dry milk and 2% normal goat serum in DPBS containing 0.05% Tween-20 (DPBS-T) for 1 h. Purified mAbs were diluted serially in DPBS-T (pH 7.4), or DPBS-T that was adjusted to pH 5.5 or 4.5 with hydrochloric acid, added to the wells and incubated for 1 h at ambient temperature. The bound antibodies were detected using goat anti-human IgG conjugated with horseradish peroxidase (Southern Biotech) diluted in blocking buffer and TMB substrate (ThermoFisher Scientific). Color development was monitored, 1N hydrochloric acid was added to stop the reaction, and the absorbance was measured at 450 nm using a spectrophotometer (Biotek).

#### Epitope mapping using an EBOV GP alanine-scan mutation library

Epitope mapping was carried out as described previously (Gilchuk et al., 2018). Comprehensive alanine scanning (‘shotgun mutagenesis’) was carried out on an expression construct for EBOV GP (Yambuku-Mayinga variant) lacking the mucin-like domain (residues 311-461), mutagenizing GP residues 33-310 and 462-676 to create a library of clones, each representing an individual point mutant. Residues were changed to alanine (with alanine residues changed to serine). The resulting library, covering 492 of 493 (99.9%) of target residues, was arrayed into 384-well plates, one mutant per well, then transfected into HEK-293T cells and allowed to express for 22 hrs. Cells, unfixed or fixed in 4% paraformaldehyde, were incubated with primary antibody and then with an Alexa Fluor 488-conjugated secondary antibody (Jackson ImmunoResearch Laboratories). After washing, cellular fluorescence was detected using the Intellicyt flow cytometer. MAb reactivity against each mutant EBOV GP clone was calculated relative to wild-type EBOV GP reactivity by subtracting the signal from mock-transfected controls and normalizing to the signal from wild-type GP-transfected controls. Mutated residues within clones were identified as critical to the mAb epitope if they did not support reactivity of the test mAb but did support reactivity of other control EBOV mAbs. This counter-screen strategy facilitated the exclusion of GP mutants that were misfolded locally or that exhibited an expression defect. The detailed algorithms used to interpret shotgun mutagenesis data were described previously (Davidson and Doranz, 2014).

#### Mouse challenge and protection studies

Groups of 7-8-week-old female BALB/c mice (n = 5 per group) housed in microisolator cages were inoculated with 1,000 PFU of the EBOV-MA by the intraperitoneal (i.p.) route. Mice were treated i.p. with 100 μg (∼5 mg/kg) of individual mAb per mouse on 1 dpi. Human mAb 2D22 (specific to an unrelated target, dengue virus) served as a negative control (Fibriansah and Lok, 2016). Mice were monitored twice daily from day 0 to 14 post infection for illness, survival, and weight loss, followed by once daily monitoring from 15 dpi to the end of the study at 28 dpi. The extent of disease was scored using the following parameters: dyspnea (possible scores 0–5), recumbence (0–5), unresponsiveness (0–5), and bleeding/hemorrhage (0–5). Moribund mice were euthanized as per the IACUC-approved protocol. All mice were euthanized on day 28 after EBOV challenge.

#### Cryo-EM trimer stability assay

Complexes for trimer stability assays were derived from data collected for structural evaluation (*see Cryo-EM sample preparation* section below). Particle picks were completed using a difference of gaussian method with low thresholds in order to pick everything on the grids. Particles were separated into stacks for either intact particles or particles that were falling apart, which was judged by eye, and then counted to determine approximate percentage of glycan cap antibody-induced instability. We have previously determined that base binding antibodies alone do not induce trimer instability.

#### Cell surface displayed GP mAb competition-binding

Jurkat-EBOV GP_CL_ cells were pre-incubated with a saturating concentration (typically 20 μg/mL) of glycan cap mAbs at room temperature for 30 min, followed by addition of labeled antibody MR78 (Flyak et al., 2015; Hashiguchi et al., 2015) at 5 μg/mL and incubated for an additional 30 min. Antibody MR78 was labeled with Alexa Fluor 647 and added after the first mAb and without washing of cells to minimize a dissociation of the first mAb from cell surface GP during a prolonged incubation. Cells were washed, fixed with 4% paraformaldehyde, and cell staining was analyzed using an iQue Screener Plus flow cytometer flow cytometer. Background values were determined from binding of the second labeled mAbs to untransfected Jurkat. Results are expressed as the percent of binding in the presence of glycan cap mAb relative to MR78 alone (maximal binding) minus background. The antibodies were considered competing if the presence of first antibody reduced the signal of the second antibody to less than 35% of its maximal binding or non-competing if the signal was greater than 86%. A level of 36–85% was considered partial competition. Thermolysin cleavage removes the epitope for most tested glycan cap antibodies that showed very low binding to Jurkat-EBOV GP_CL_ (data not shown). This study served as an additional control to confirm that cleavage inhibition measured as percent of RBS exposure is not due to MR78 binding completion with residually bound glycan cap antibody on Jurkat-EBOV GP_CL_.

#### Cell surface displayed GP cleavage inhibition

Jurkat-EBOV GP cells were pre-incubated with serial dilutions of mAbs in PBS for 20 min at room temperature, then incubated with thermolysin (Promega) for 20 min at 37°C. The reaction was stopped by addition of the incubation buffer containing DPBS, 2% heat inactivated FBS and 2 mM EDTA (pH 8.0). Washed cells were incubated with 5 μg/mL of fluorescently labeled RBS-specific mAb MR78 at 4°C for 60 min. Stained cells were washed, fixed, and analyzed by flow cytometry using iQue Screener Plus flow cytometer. Cells were gated for the viable population. Background staining was determined from binding of the labeled mAb MR78 to Jurkat-EBOV GP (uncleaved) cells. Results are expressed as the percent of RBS exposure in the presence of tested mAb relative to labeled MR78 mAb-only control (maximal binding to Jurkat-EBOV GP_CL_) minus background. The GP base-directed antibody 2G4 (Qiu et al., 2011) and 2D22 served as negative controls. BDBV-329 was excluded because it does not bind to EBOV and BDBV-43 was excluded due to poor recombinant expression of the antibody.

#### Construct design, expression and protein purification

EBOV GP (Makona) (residues 32-644, GenBank AKG65268.1) with an N-terminal tissue plasminogen activator (*Homo sapiens*) signal sequence was codon optimized for mammalian protein expression, synthesized and subcloned into the pPPI4 expression vector (GenScript). A C-terminal enterokinase (Ek) cut site (DDDDK) was introduced after residue 628 followed by a short linker (AG) and two streptavidin tags (WSHPQFEK) separated by a GS-linker (GGGSGGGSGGGS). Residues 310-460 were removed to produce EBOV GPΔMuc. BDBV GP (residues 1-643, GenBank ALT19772.1) with the GP-associated signal peptide, an Ek cut site after residue 643 followed by an AG-linker and the double streptavidin tag as described above was codon optimized for mammalian protein expression, synthesized and subcloned into pPPI4. Residues 313-470 were removed to produce BDBV GPΔMuc. EBOV sGP (Mayinga) (residues 1-314, GenBank AAD14584.1) with the sGP-associated signal peptide an enterokinase cut site after residue 314 followed by an AG-linker and a double streptavidin tag was codon optimized for mammalian protein expression, synthesized and subcloned into pPPI4.

All GPs were expressed and purified in transiently transfected HEK-293F cells at a density of 0.8-1.5 x 10^6^ cells/mL using 750 μg of DNA and 2.25 mg of polyethylenimine “Max” (MW 25,000, Polyscience, Inc.) mixed with 50 mL of Opti-MEM (ThermoFisher Scientific). Solutions were sterile filtered using 0.22 μm Steriflip disposable filters (Millipore) and allowed to incubate at room temperature for 30 min before being added to cultures. After 5 days of expression at 37°C and 8% CO_2_, cells were harvested by centrifugation (8,000 x g for 1hr at 4°C) and filtered to remove cellular debris. BioLock biotin blocking solution (IBA Lifesciences) was added to supernatant according to the manufacturer’s protocol before being loaded onto Strep-Tactin Superflow Plus beads (Qiagen) that had been pre-equilibrated in 1X Strep Buffer (100 mM Tris, pH 8.0, 150 mM NaCl and 1mM EDTA). Beads were washed with 10 mL of 1X Strep Buffer and GPs were eluted by addition of 2.5 mM d-desthiobiotin added to 10 mL of 1X Strep Buffer. GPs were further purified by size exclusion chromatography (SEC) using an S200 increase (S200I, GE) column equilibrated in 1X TBS (150 mM NaCl, 20 mM Tris, pH 7.4).

For EBOV-237, BDBV-329, EBOV-442, EBOV-437 and 2D22 recombinant mAb production, cDNA encoding the genes of heavy and light chains were cloned into the pTwist CMV Betaglobin WPRE Neo vector encoding IgG1 or Fab-heavy chain (McLean et al., 2000), or monocistronic expression vector pTwist-mCis_G1 (Zost et al., 2020) and transformed into E. coli cells. mAb proteins were produced after transient transfection of ExpiCHO cells following the manufacturer’s protocol and were purified from filtered culture supernatants by fast protein liquid chromatography (FPLC) on an AKTA instrument using HiTrap MabSelect Sure column for IgG (GE Healthcare Life Sciences) or CaptureSelect™ IgG-CH1 column for Fab (ThermoFisher Scientific). Purified mAbs were buffer exchanged into PBS, filtered using sterile 0.45 μm pore size filter devices (Millipore), concentrated, and stored in aliquots at −80°C until use.

For BDBV-289, BDBV-43, EBOV-293 and EBOV-296 antibody expression, sequences were optimized for mammalian expression, synthesized and subcloned into the expression vector AbVec containing the human IgG HC constant region or the human lambda or kappa LC constant region (GenScript). Fab was produced by the introduction of a stop codon after residue 226 in the HC hinge-region. ADI-15878 Fab and ADI-16061 Fab were used as a fiducials in this study and were produced as previously described (Murin et al., 2018). IgGs and Fab were transiently transfected as described above for GPs, except that 500 μg of HC DNA and 250 μg of LC DNA was mixed to encourage HC/LC pairing and the avoidance of LC dimers. For BDBV-289 and BDBV-43 Fab, cell supernatants were loaded onto 5 mL Lambda (BDBV-289) or Kappa (BDBV-43) Select columns (GE) that had been equilibrated in 1X phosphate buffered saline (PBS, QualityBiological) followed by elution with 0.1 M glycine, pH 3.0. Fab were subsequently buffer exchanged into 20 mM sodium acetate (NaOAc), pH 5.6 by dialysis and loaded onto a MonoS column (GE). Fab were then eluted with a gradient of 1M KCl. For EBOV-437, EBOV-442, EBOV-293 and EBOV-296 Fab, cell supernatants were loaded onto a 1 mL or 5 mL Capture Select column (ThermoFisher Scientific) and eluted with 0.1 M glycine, pH 3.0. Appropriate fractions were pooled and further purified by SEC using an S200I column equilibrated in 1X TBS buffer. For IgG, supernatants were loaded onto a HiTrap 5 mL mAb Select column (GE) that had been pre-equilibrated in 1X PBS followed by elution with 0.1 M glycine, pH 3.0 and neutralization with 1M Tris, pH 8.5. Appropriate fractions were pooled and further purified by SEC using an S200I column that had been equilibrated with 1X TBS.

#### Crystallization and Structure Determination of BDBV-289 Fab

Fabs produced for crystallographic studies were made in Expi-CHO cells per the manufacturer’s “max titer” protocol (GIBCO/ThermoFisher Scientific) and purified as described above. BDBV-289 Fab was screened for crystallization using the Joint Center for Structural Genomics (JCSG) Rigaku CrystalMation at The Scripps Research Institute against the JCSG Core Suites I-IV. Protein at 7.4 mg/mL was mixed 1:1 with precipitants and crystallized using the vapor diffusion method at both room temperature and 4°C. Crystals grew in 0.1M HEPES pH 6.5 and 20% (w/v) polyethylene glycol 6000 at 4°C. Crystals were cryoprotected with well solution augmented with 30% ethylene glycol. Data were collected at Stanford Synchrotron Radiation Light Source beamline 12-2. Data were indexed, integrated and scaled using HKL-2000 (Otwinowski and Minor, 1997) to 3.0 Å (Table S2). Crystals belonged to the space group P6_1_ with a single Fab in the asymmetric unit.

Data were phased using Phaser (McCoy et al., 2007) with molecular replacement by a homology model generated using Swiss Modeler (Biasini et al., 2014). A single Fab was placed in the asymmetric unit. This initial solution was rebuilt manually in Coot (Emsley et al., 2010), followed by multiple rounds of refinement in Phenex.refine (Adams et al., 2010) and model building with Coot. Translation/Libration/Screw (TLS) groups were introduced towards the end of refinement. Four TLS groups were set manually with one for each immunoglobulin domain. A large positive density seen in the difference map was modeled as PEG after evaluating fits for all components of the buffer.

#### Cryo-EM sample preparation

EBOV/Mak GPΔmuc was incubated overnight with a 5-fold molar excess of each Fab at 4°C. The complexes were then purified by SEC using an S200I column equilibrated in 1X TBS and concentrated using a 100-kDa concentrator (Amicon Ultra, Millipore) and mixed with detergent immediately prior to freezing (**Table S2**). Vitrification was performed with a Vitrobot (ThermoFisher Scientific) equilibrated to 4°C and 100% humidity. Cryo-EM grids were plasma cleaned for 5s using a mixture of Ar/O_2_ (Gatan Solarus 950 Plasma system) followed by blotting on both sides of the grid with filter paper (Whatman No. 1). See Table S1 for additional details for individual complexes. Note that ADI-15878 Fab was added to the BDBV-43 complex and ADI-16061 Fab and EBOV-515 Fab was added to the EBOV-437, EBOV-442, EBOV-293, EBOV-296, EBOV-237 and BDBV-329 complexes to assist in angular sampling and orientations in ice.

#### Cryo-EM data collection and processing

Cryo-EM data were collected according to Table S1. Micrographs were aligned and dose-weighted using MotionCor2 (Zheng et al., 2017). CTF estimation was completed using GCTF (Zhang, 2016). Particle picking and initial 2D classification were initially performed using CryoSPARC 2.0 (Punjani et al., 2017) to clean up particle stacks and exclude any complexes that were degrading. For those reconstructions that required more extensive 3D classification, particle picks were then imported into Relion 3.1b1 (Zivanov et al., 2018) for 3D classification and then refinement using appropriate symmetry where necessary and a tight mask around the GP/Fab complex of interest. CTF refinement was then performed by either Relion or CryoSPARC to increase map quality and resolution. There was no density for ADI-16061 in any of the maps and we did not build density for ADI-15878 in the BDBV-43 map (this was previously deposited under PDB 6DZM). We chose not to build a model into EBOV-515 density but included this density in our reconstructions to assist with particle alignment.

#### Cryo-EM model building and refinement

Homology models of Fab were first generated using SWISS-MODEL (Biasini et al., 2014). Models of BDBV GP (PDB: 6DZM) and EBOV GP (PDB: 5JQ3) were then added to generate starting models used for refinement. Models were fit into maps using UCSF Chimera (Pettersen et al., 2004) and refined initially using Phenix real-space refinement using NCS constraints (Liebschner et al., 2019). The refined model was then used as a template for relaxed refinement in Rosetta (DiMaio et al., 2015). The top five models were then evaluated for fit in EM density and adjusted manually using Coot (Emsley et al., 2010) to maximize fit. Finally, Man9 glycans were fit into glycan densities, trimmed and then a final refinement was performed in Rosetta. The final structures were evaluated using EMRinger (Barad et al., 2015) and Molprobity from Phenix. All figures were generated in UCSF Chimera (Pettersen et al., 2004). Antibody contacts were analyzed using LigPlot (Laskowski and Swindells, 2011), Arpeggio (Jubb et al., 2017) and UCSF Chimera (Pettersen et al., 2004).

#### Inferred germline antibody analysis

Inferred germline sequences for BDBV289 and BDBV43 F_V_ domains were determined using IMGT/V-QUEST (Brochet et al., 2008; Giudicelli et al., 2011). Nucleotide sequences of B-cells originally isolated from donors were kindly provided by James Crowe and used to derive a list of likely germline VDJ genes. Those with the highest confidence were then used to reconstruct an inferred germline sequence. The mature CDRH3 sequence was included in the reconstructed germline sequences due to low confidence in predicting germline CDRH3 sequences, although some residues were predicted to be different from the germline CDRH3. For BDBV-289 and BDBV-43, inferred germline sequences were then codon optimized for mammalian protein expression and sub-cloned into the appropriate AbVec expression vector. Stop codons were introduced as described above to produce Fab.

### QUANTIFICATION AND STATISTICAL ANALYSIS

The descriptive statistics mean ± SEM or mean ± SD were determined for continuous variables as noted. EC_50_ values for mAb binding were determined after log transformation of antibody concentration using sigmoidal dose-response nonlinear regression analysis. Correlation between antibody synergy and percent monomer in GP trimer fraction was estimated using linear regression analysis. In neutralization assays, IC_50_ values were calculated after log transformation of antibody concentrations using a 4-parameter nonlinear fit analysis. Technical and biological replicates are indicated in the figure legends. Statistical analyses were performed using Prism v8 (GraphPad).

