## Supplemental Materials for "Convergence of a common solution to broad ebolavirus neutralization by glycan cap directed human antibodies"

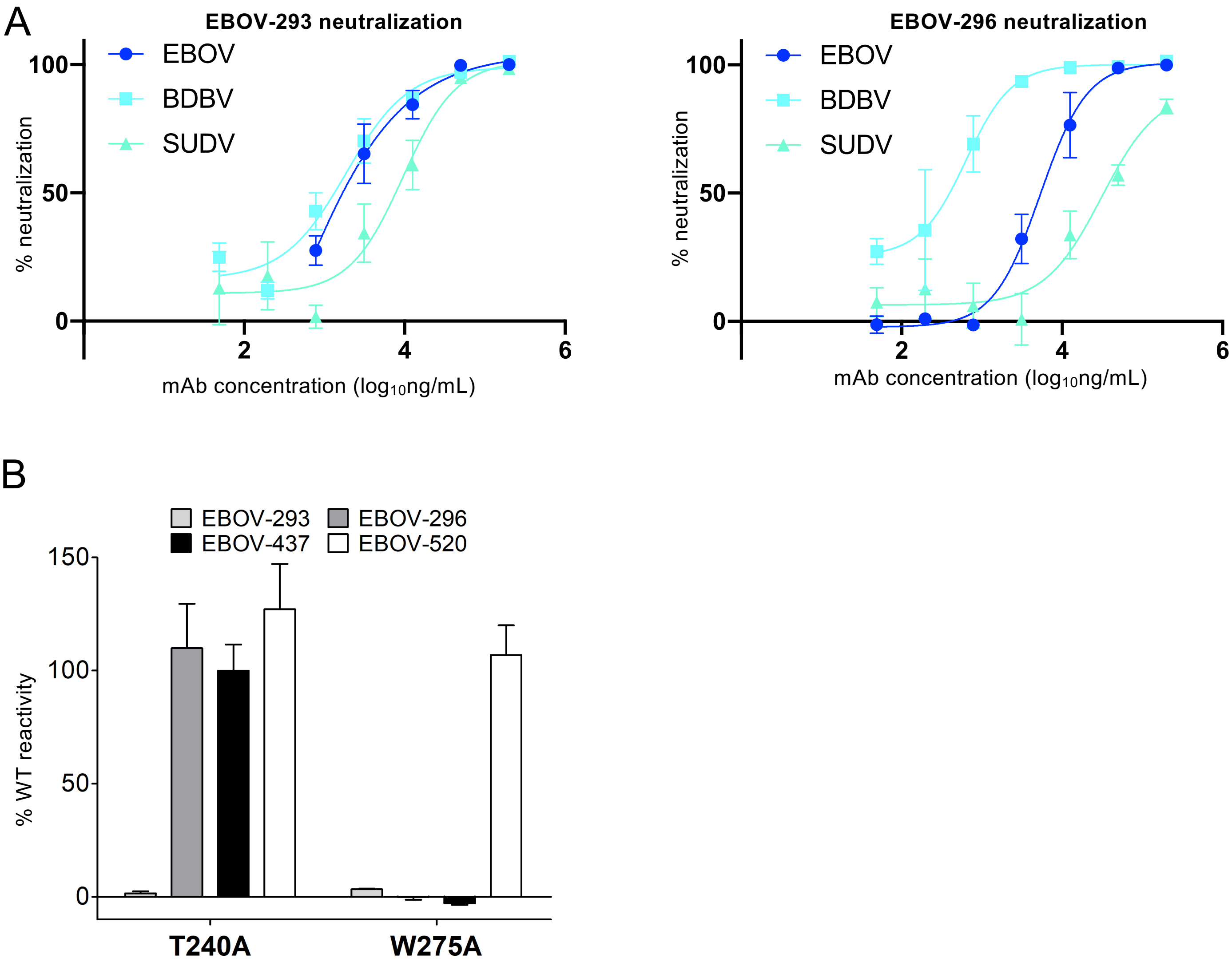

A
EBOV-293 neutralization
EBOV
BDBV
SUDV
% neutralization
100
50
0
2
4
6
mAb concentration (log10ng/mL)
EBOV-296 neutralization
EBOV
BDBV
SUDV
% neutralization
100
50
0
2
4
6
mAb concentration (log10ng/mL)
B
EBOV-293
EBOV-296
EBOV-437
EBOV-520
% WT reactivity
150
100
50
0
T240A
W275A

**Figure S1. Characterization of EBOV-293 and EBOV-296 antibodies from a human individual, related to Table 1.** **(A)** Neutralizing data for EBOV-293 and EBOV-293 (hybridoma-derived antibodies) for the three major ebolaviruses. $IC_{50}$ values are listed in Table 1. **(B)** Site-directed mutagenesis of residues determined by alanine scanning to be important for the binding of EBOV-293 and EBOV-296. EBOV-437 and EBOV-520 are included for reference and as controls.

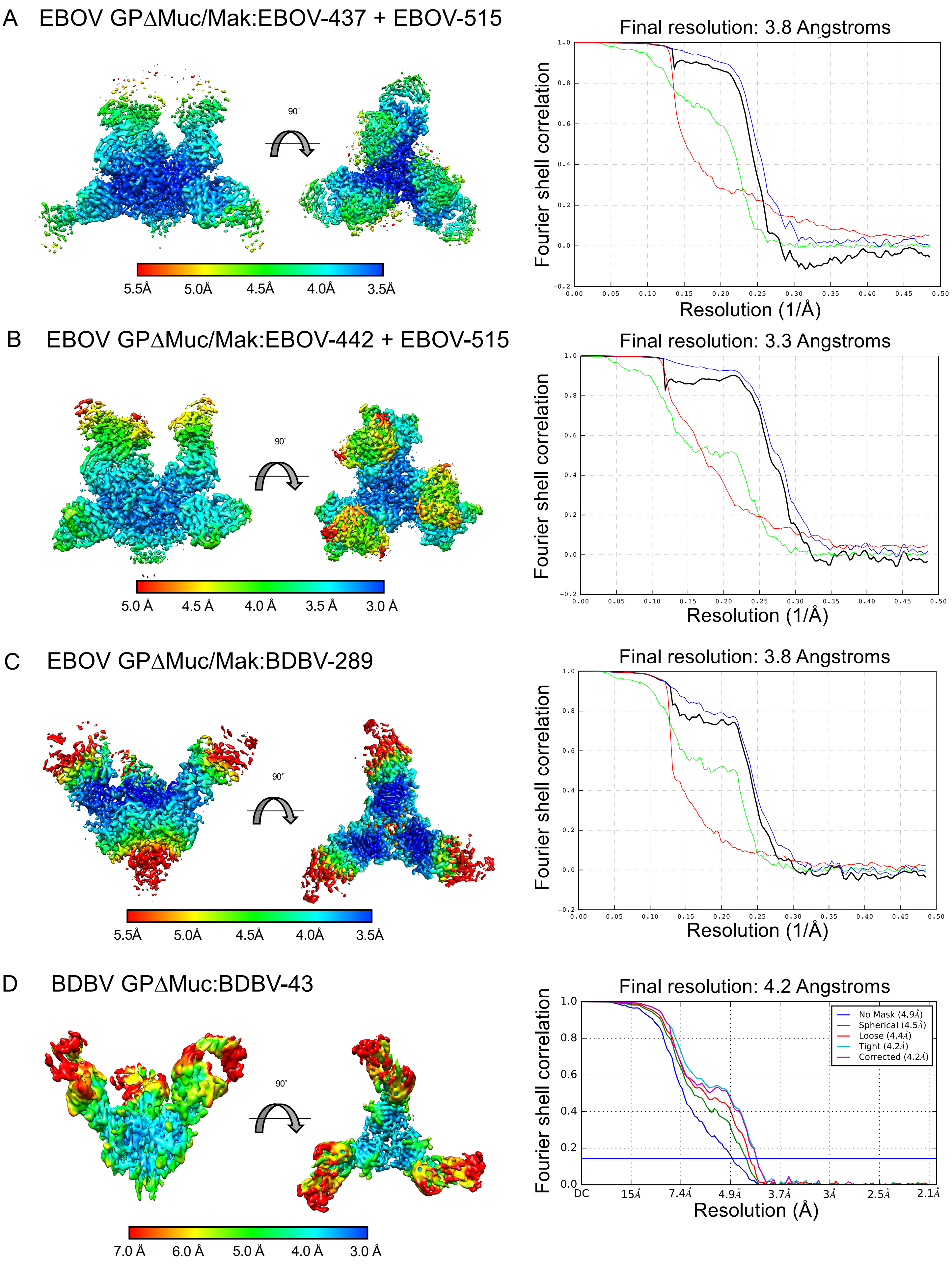
A EBOV GPΔMuc/Mak:EBOV-437 + EBOV-515
90°
5.5Å 5.0Å 4.5Å 4.0Å 3.5Å
Final resolution: 3.8 Angstroms
Fourier shell correlation
Resolution (1/Å)
B EBOV GPΔMuc/Mak:EBOV-442 + EBOV-515
90°
5.0 Å 4.5 Å 4.0 Å 3.5 Å 3.0 Å
Final resolution: 3.3 Angstroms
Fourier shell correlation
Resolution (1/Å)
C EBOV GPΔMuc/Mak:BDBV-289
90°
5.5Å 5.0Å 4.5Å 4.0Å 3.5Å
Final resolution: 3.8 Angstroms
Fourier shell correlation
Resolution (1/Å)
D BDBV GPΔMuc:BDBV-43
90°
7.0 Å 6.0 Å 5.0 Å 4.0 Å 3.0 Å
Final resolution: 4.2 Angstroms
No Mask (4.9Å)
Spherical (4.5Å)
Loose (4.4Å)
Tight (4.2Å)
Corrected (4.2Å)
Fourier shell correlation
DC 15Å 7.4Å 4.9Å 3.7Å 3Å 2.5Å 2.1Å
Resolution (Å)

E EBOV GPΔMuc/Mak:EBOV-293 + EBOV-515

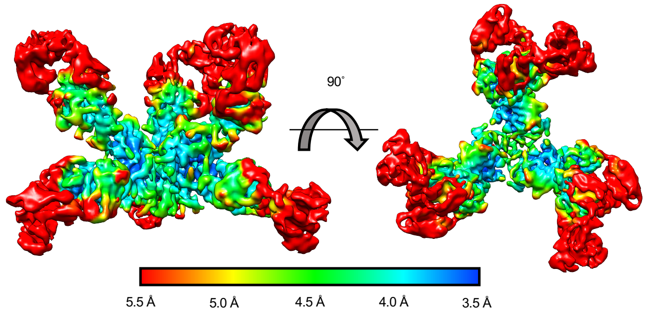


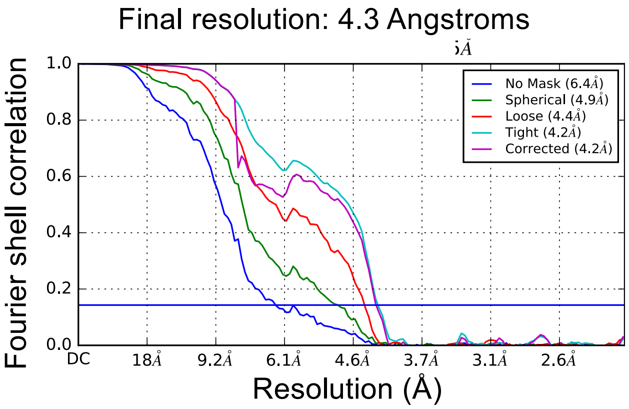


F EBOV GPΔMuc/Mak:EBOV-296 + EBOV-515

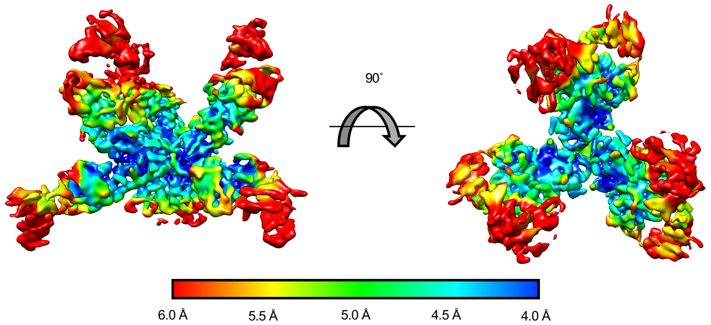


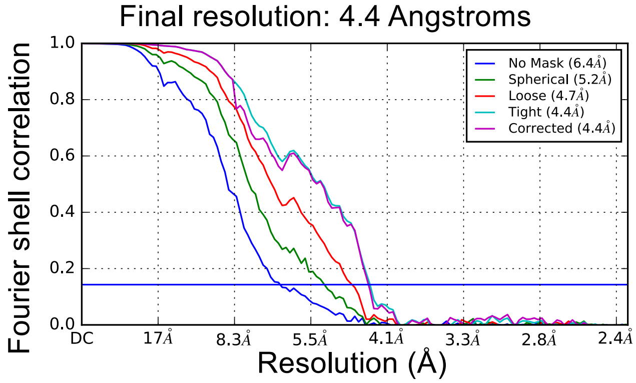


G BDBV GPΔMuc/Mak:BDBV-329 + EBOV-515

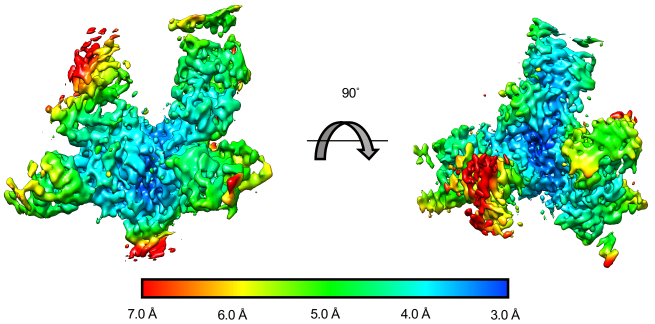


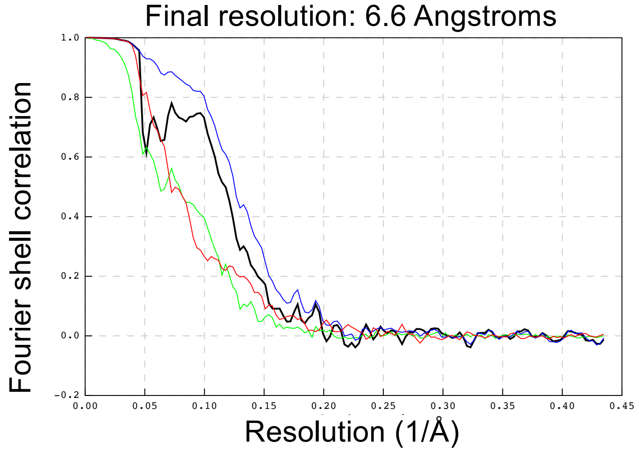


H EBOV GPΔMuc/Mak:EBOV-237 + EBOV-515

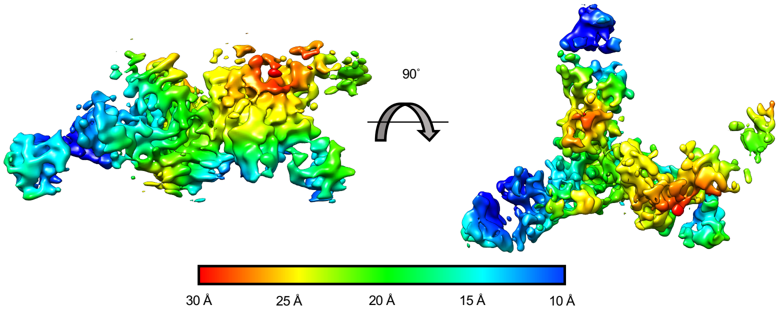


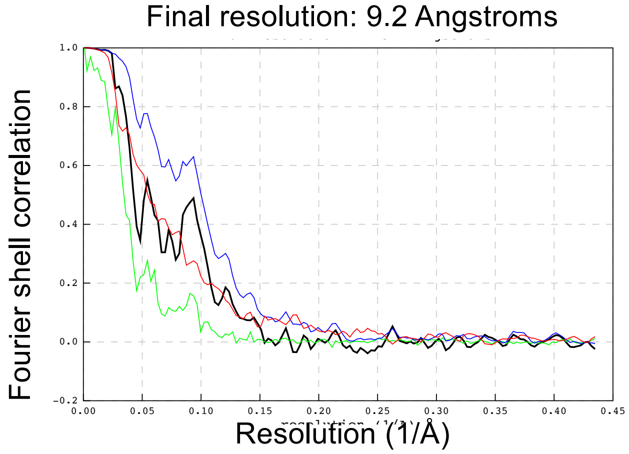

**Figure S2. Cryo-EM local resolution estimation and FSC curves, related to Figure 2.** Local resolution is plotted onto the cryo-EM maps of structures solved in this study according to the scale for each figure. Side views (left) and top views (right) are shown, with respect to the viral membrane. Fourier shell correlation curves for each reconstruction are shown with the estimated resolution indicated. Relion 3.1 was used in panels A to C and G to H. Cryosparc 2.0 was used for reconstructions in panels D to F.

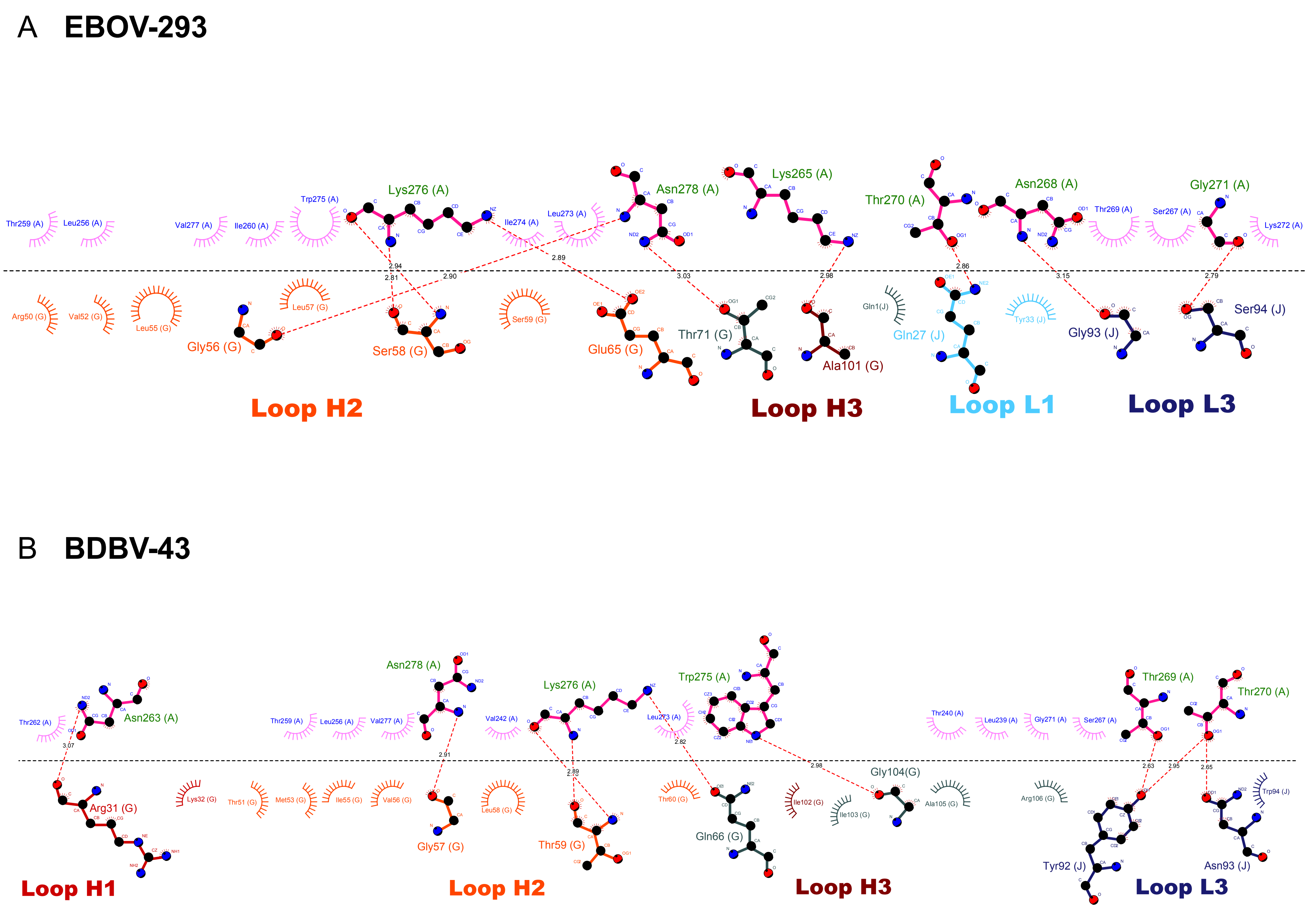

A EBOV-293
Loop H2
Loop H3
Loop L1
Loop L3
B BDBV-43
Loop H1
Loop H2
Loop H3
Loop L3

C **EBOV-437**

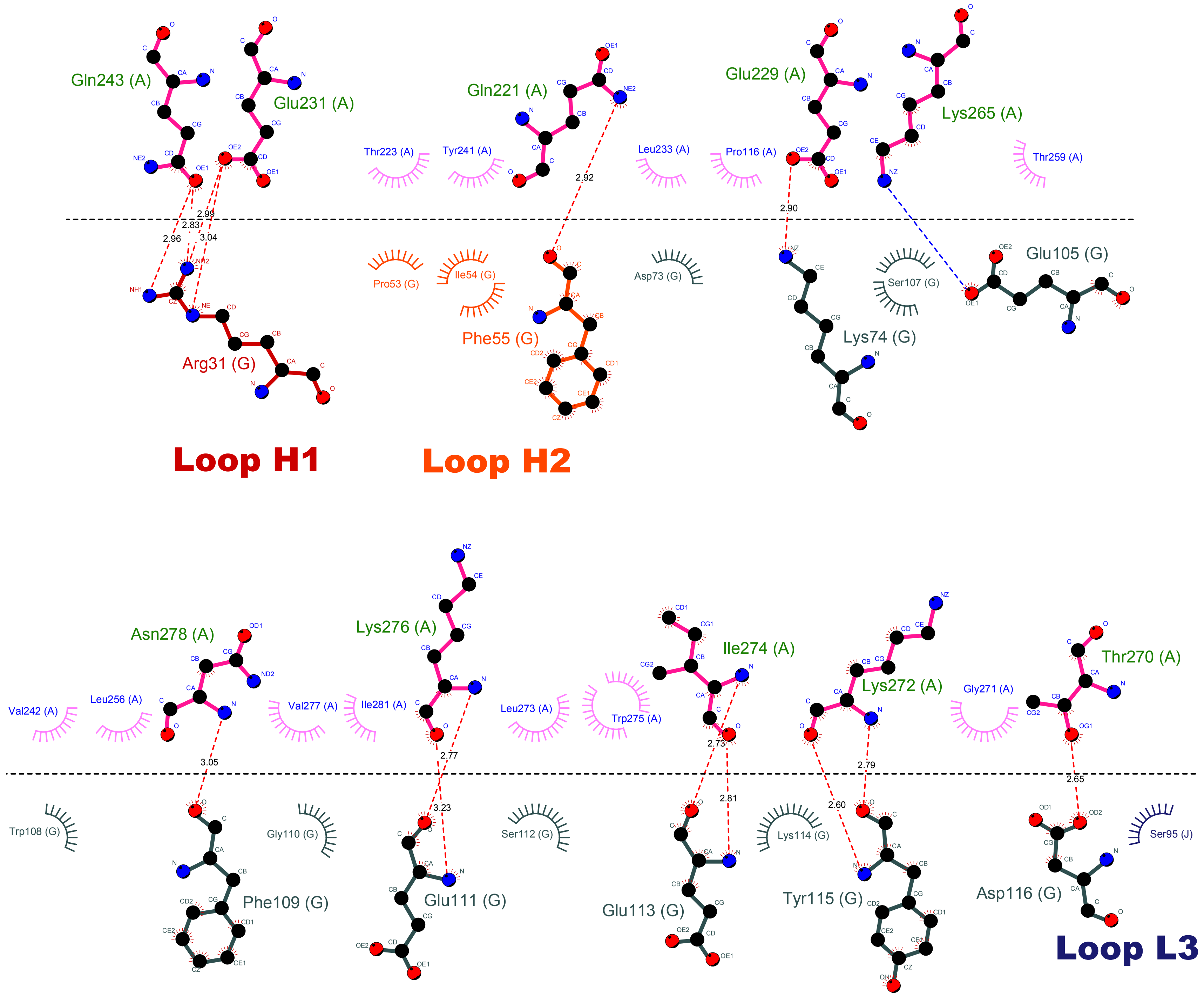

Gln243 (A)
Glu231 (A)
Thr223 (A)
Tyr241 (A)
Gln221 (A)
Leu233 (A)
Pro116 (A)
Glu229 (A)
Lys265 (A)
Thr259 (A)
2.99
2.83
2.96
3.04
2.92
2.90
Pro53 (G)
Ile54 (G)
Asp73 (G)
Ser107 (G)
Glu105 (G)
Arg31 (G)
Phe55 (G)
Lys74 (G)
Loop H1
Loop H2
Asn278 (A)
Lys276 (A)
Ile274 (A)
Lys272 (A)
Thr270 (A)
Val242 (A)
Leu256 (A)
Val277 (A)
Ile281 (A)
Leu273 (A)
Trp275 (A)
Gly271 (A)
3.05
2.77
3.23
2.73
2.81
2.60
2.79
2.65
Trp108 (G)
Gly110 (G)
Ser112 (G)
Lys114 (G)
Ser95 (J)
Phe109 (G)
Glu111 (G)
Glu113 (G)
Tyr115 (G)
Asp116 (G)
Loop L3

D **BDBV-289**

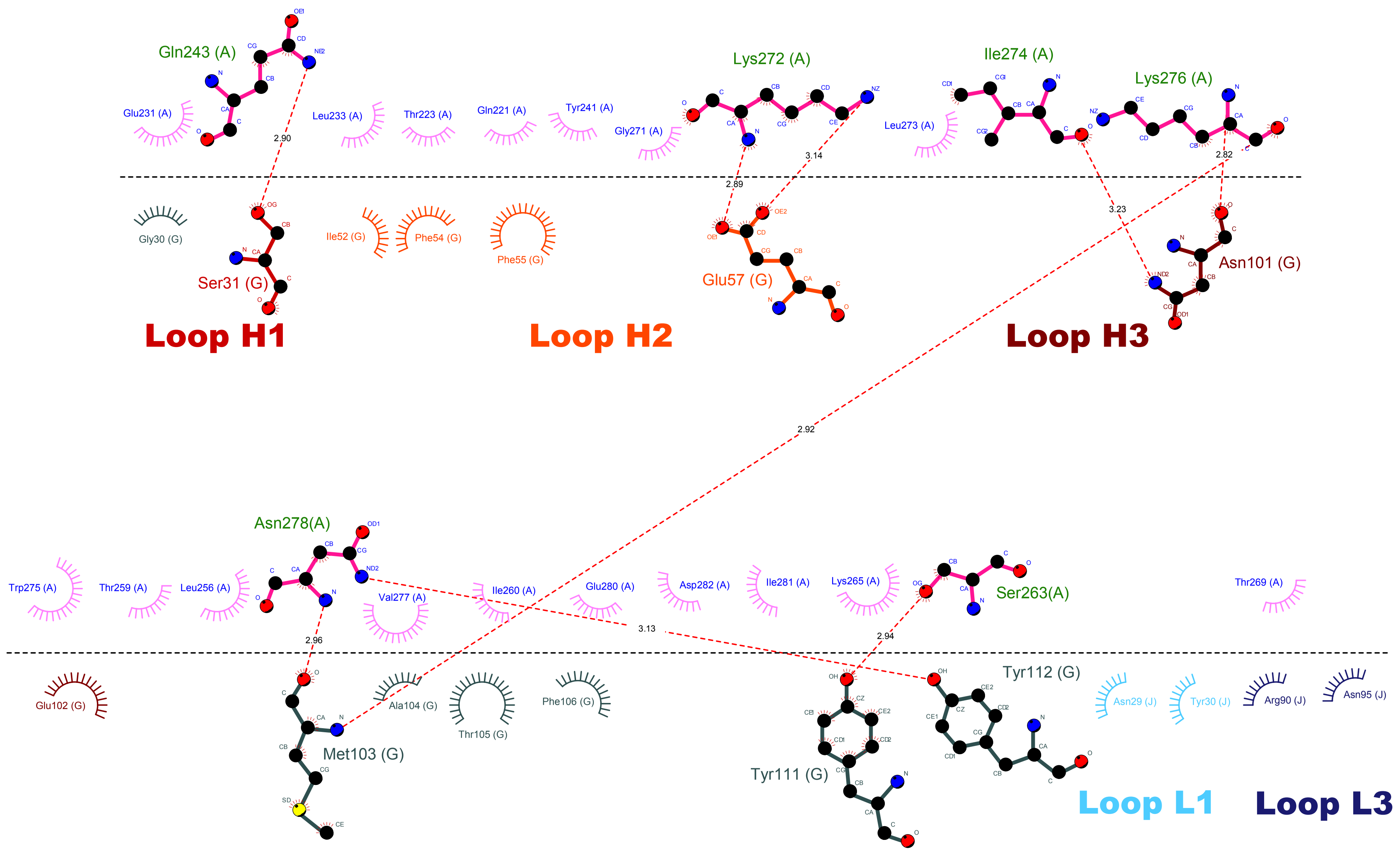

Gln243 (A)
Glu231 (A)
Leu233 (A)
Thr223 (A)
Gln221 (A)
Tyr241 (A)
Gly271 (A)
Lys272 (A)
Ile274 (A)
Lys276 (A)
Leu273 (A)
2.90
2.89
3.14
3.23
2.82
Gly30 (G)
Ser31 (G)
Ile52 (G)
Phe54 (G)
Phe55 (G)
Glu57 (G)
Asn101 (G)
Loop H1
Loop H2
Loop H3
2.92
Asn278(A)
Trp275 (A)
Thr259 (A)
Leu256 (A)
Val277 (A)
Ile260 (A)
Glu280 (A)
Asp282 (A)
Ile281 (A)
Lys265 (A)
Ser263(A)
Thr269 (A)
2.96
3.13
2.94
Glu102 (G)
Met103 (G)
Ala104 (G)
Thr105 (G)
Phe106 (G)
Tyr111 (G)
Tyr112 (G)
Asn29 (J)
Tyr30 (J)
Arg90 (J)
Asn95 (J)
Loop L1
Loop L3

E EBOV-442

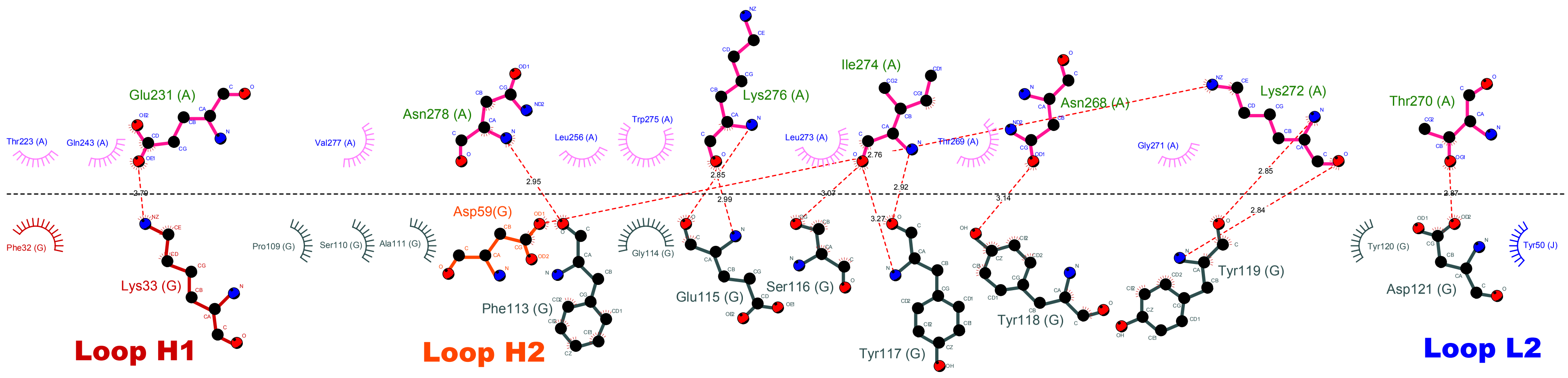

Glu231 (A)
Asn278 (A)
Lys276 (A)
Ile274 (A)
Asn268 (A)
Lys272 (A)
Thr270 (A)
Thr223 (A)
Gln243 (A)
Val277 (A)
Leu256 (A)
Trp275 (A)
Leu273 (A)
Thr269 (A)
Gly271 (A)
2.79
2.95
2.85
2.99
3.07
2.76
2.92
3.27
3.14
2.85
2.84
2.87
Phe32 (G)
Lys33 (G)
Pro109 (G)
Ser110 (G)
Ala111 (G)
Asp59(G)
Phe113 (G)
Gly114 (G)
Glu115 (G)
Ser116 (G)
Tyr117 (G)
Tyr118 (G)
Tyr119 (G)
Tyr120 (G)
Asp121 (G)
Tyr50 (J)
Loop H1
Loop H2
Loop L2

F EBOV-296

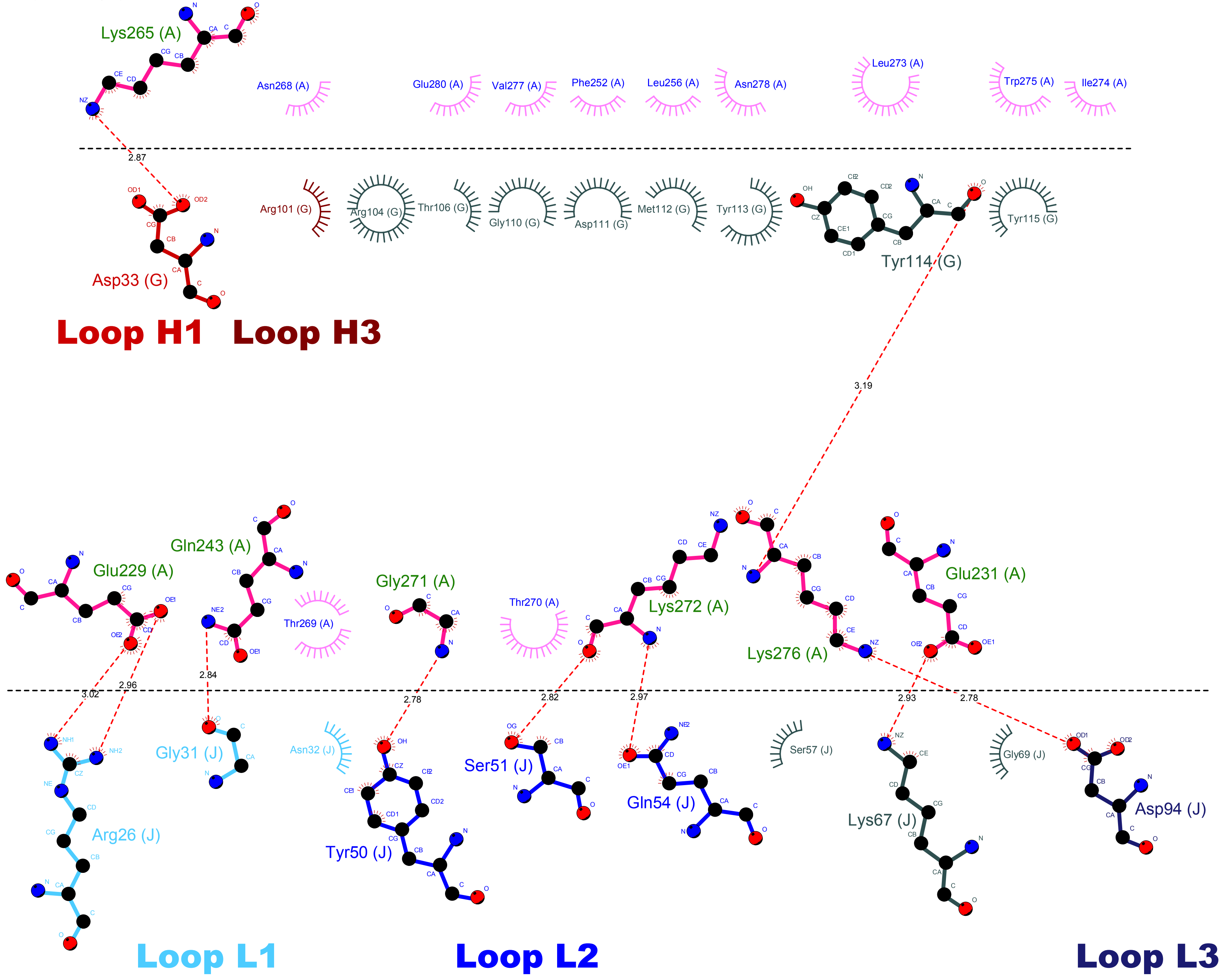

Lys265 (A)
Asn268 (A)
Glu280 (A)
Val277 (A)
Phe252 (A)
Leu256 (A)
Asn278 (A)
Leu273 (A)
Trp275 (A)
Ile274 (A)
2.87
Asp33 (G)
Arg101 (G)
Arg104 (G)
Thr106 (G)
Gly110 (G)
Asp111 (G)
Met112 (G)
Tyr113 (G)
Tyr114 (G)
Tyr115 (G)
Loop H1
Loop H3
3.19
Glu229 (A)
Gln243 (A)
Thr269 (A)
Gly271 (A)
Thr270 (A)
Lys272 (A)
Lys276 (A)
Glu231 (A)
3.02
2.96
2.84
2.78
2.82
2.97
2.93
2.78
Arg26 (J)
Gly31 (J)
Asn32 (J)
Tyr50 (J)
Ser51 (J)
Gln54 (J)
Ser57 (J)
Lys67 (J)
Gly69 (J)
Asp94 (J)
Loop L1
Loop L2
Loop L3

**Figure S3. Glycan cap antibody epitope and paratope interactions, related to Figure 3.** LigPlot schematics **(A to E)** for each of the glycan cap antibody structures solved in this study are shown. Plots are labeled according to the description in Fig. S3. Related to Figures 4 and 5.

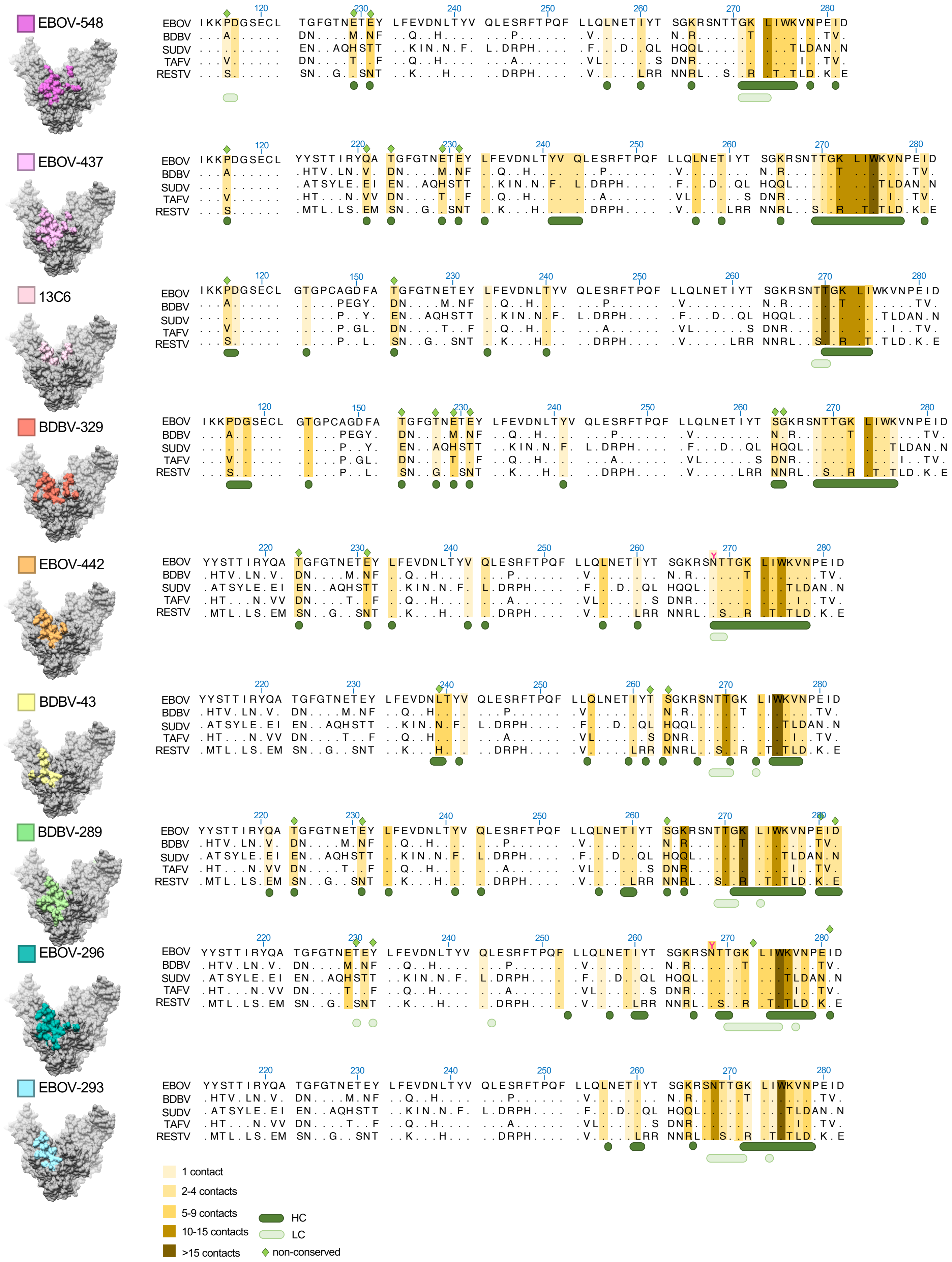
EBOV-548
EBOV-437
13C6
BDBV-329
EBOV-442
BDBV-43
BDBV-289
EBOV-296
EBOV-293
EBOV
BDBV
SUDV
TAFV
RESTV
1 contact
2-4 contacts
5-9 contacts
10-15 contacts
>15 contacts
HC
LC
non-conserved

**Figure S4. Epitope conservation of glycan cap antibodies, related to Figures 2 and 3.** Apo-GP is rendered in grey on the left with the corresponding glycan cap antibody footprint highlighted. On the right are aligned sequences of the interacting regions on GP. Total contacts for each residue at 4 Å distance or less were determined (see Table S3) and residues are highlighted in gold according to the number of contacts, with darker brown indicating more contacts and thus more likely to be an important residue for binding. Residues that are not well-conserved are marked with a green diamond. HC contacts are indicated below in dark green and LC contacts in light green. Related to Figure 7.

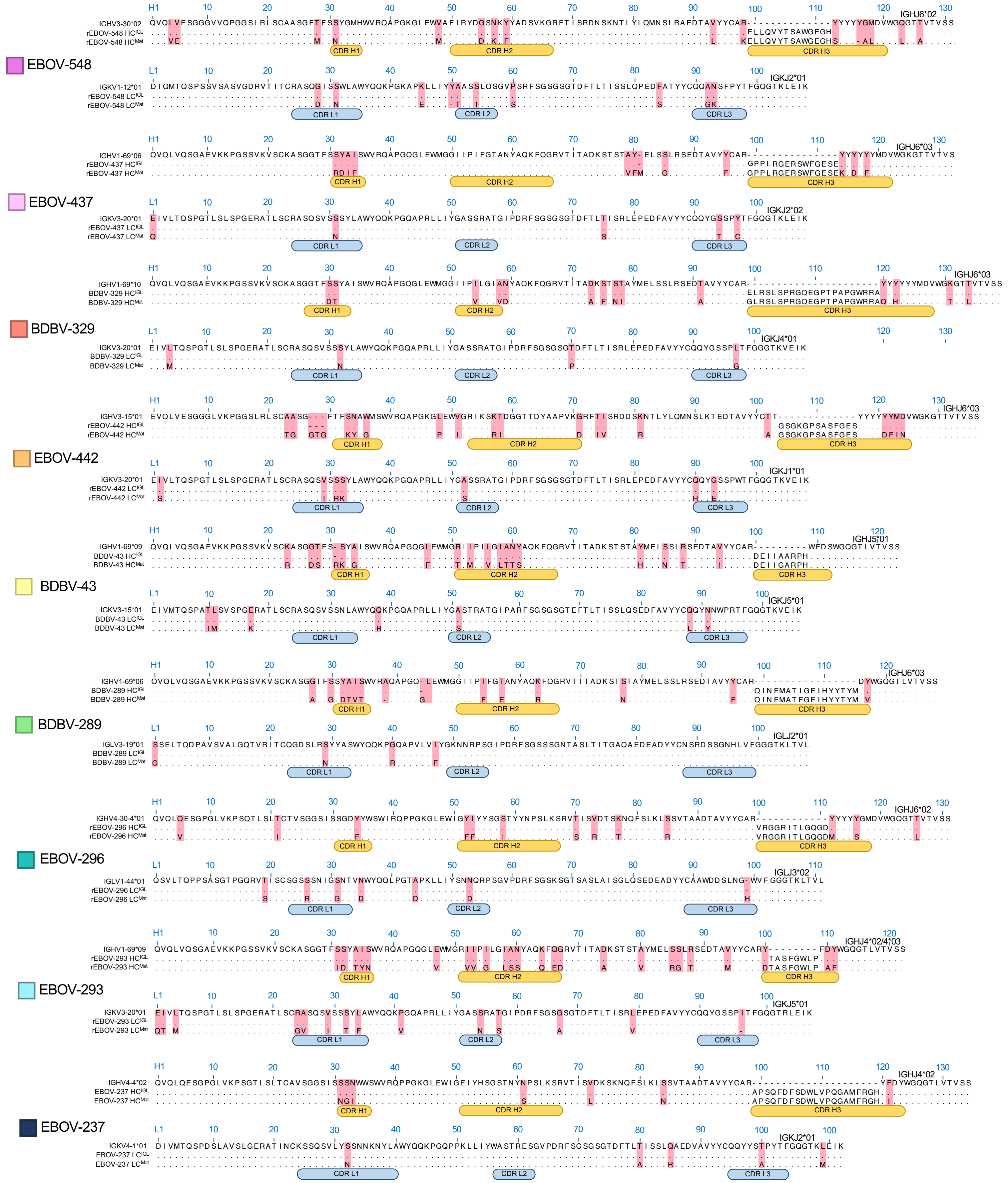

EBOV-548
IGHV3-30*02
rEBOV-548 HC^IGL
rEBOV-548 HC^Mat
IGHJ6*02
IGKV1-12*01
rEBOV-548 LC^IGL
rEBOV-548 LC^Mat
IGKJ2*01
EBOV-437
IGHV1-69*06
rEBOV-437 HC^IGL
rEBOV-437 HC^Mat
IGHJ6*03
IGKV3-20*01
rEBOV-437 LC^IGL
rEBOV-437 LC^Mat
IGKJ2*02
BDBV-329
IGHV1-69*10
BDBV-329 HC^IGL
BDBV-329 HC^Mat
IGHJ6*03
IGKV3-20*01
BDBV-329 LC^IGL
BDBV-329 LC^Mat
IGKJ4*01
EBOV-442
IGHV3-15*01
rEBOV-442 HC^IGL
rEBOV-442 HC^Mat
IGHJ6*03
IGKV3-20*01
rEBOV-442 LC^IGL
rEBOV-442 LC^Mat
IGKJ1*01
BDBV-43
IGHV1-69*09
BDBV-43 HC^IGL
BDBV-43 HC^Mat
IGHJ5*01
IGKV3-15*01
BDBV-43 LC^IGL
BDBV-43 LC^Mat
IGKJ5*01
BDBV-289
IGHV1-69*06
BDBV-289 HC^IGL
BDBV-289 HC^Mat
IGHJ6*03
IGLV3-19*01
BDBV-289 LC^IGL
BDBV-289 LC^Mat
IGLJ2*01
EBOV-296
IGHV4-30-4*01
rEBOV-296 HC^IGL
rEBOV-296 HC^Mat
IGHJ6*02
IGLV1-44*01
rEBOV-296 LC^IGL
rEBOV-296 LC^Mat
IGLJ3*02
EBOV-293
IGHV1-69*09
rEBOV-293 HC^IGL
rEBOV-293 HC^Mat
IGHJ4*02/4*03
IGKV3-20*01
rEBOV-293 LC^IGL
rEBOV-293 LC^Mat
IGKJ5*01
EBOV-237
IGHV4-4*02
EBOV-237 HC^IGL
EBOV-237 HC^Mat
IGHJ4*02
IGKV4-1*01
EBOV-237 LC^IGL
EBOV-237 LC^Mat
IGKJ2*01
CDR H1
CDR H2
CDR H3
CDR L1
CDR L2
CDR L3

**Figure S5. Inferred germline sequences of glycan cap antibodies, related to figures 4 and 5.**

The germline genes for each glycan cap antibody were determined (top line) from nucleotide sequences (see methods) and inferred germline (IGL) sequences for the HC and LC were assembled (middle line) and aligned with the mature (Mat) sequence (bottom line). HC CDR loops are indicated below in yellow and LC CDR loops are indicated in blue. Residues that changed from germline are highlighted in pink. Related to Figure 7.

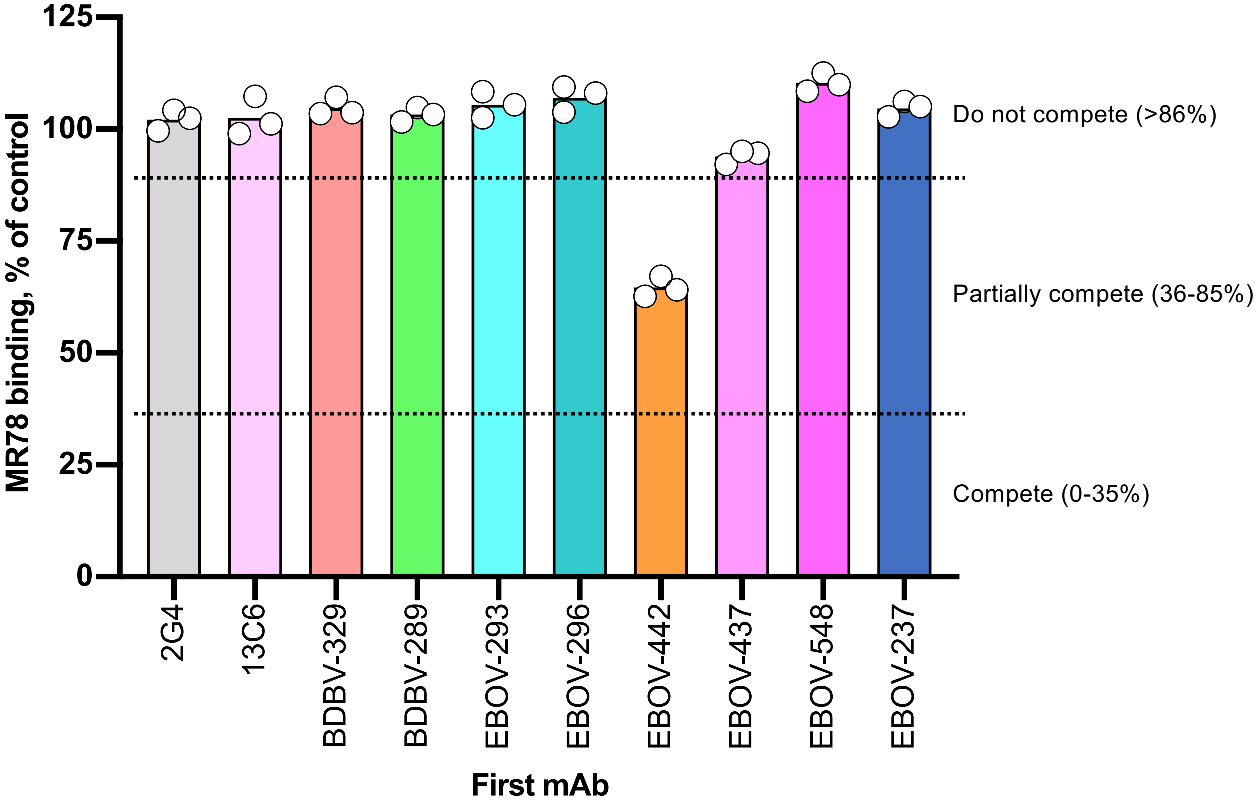
MR78 binding, % of control
125
100
75
50
25
0
Do not compete (>86%)
Partially compete (36-85%)
Compete (0-35%)
2G4
13C6
BDBV-329
BDBV-289
EBOV-293
EBOV-296
EBOV-442
EBOV-437
EBOV-548
EBOV-237
First mAb

**Figure S6**. **Control for specificity of RBS exposure measurement on Jurkat-EBOV $GP_{CL}$ in a presence of glycan cap antibody, related to Figure 6**. Jurkat-EBOV $GP_{CL}$ cells that obtained after thermolysin treatment of Jurkat-EBOV cells were pre-incubated with 20 µg/mL of respective glycan cap antibody followed by addition of fluorescently labeled antibody MR78. Binding was analyzed by flow cytometry, and the results are expressed as the percent of binding in the presence of glycan cap mAb relative to MR78 alone (maximal binding) minus background. The antibodies were considered competing if the presence of first antibody reduced the signal of the second antibody to less than 35% of its maximal binding or non-competing if the signal was greater than 86%. A level of 36–85% was considered partial competition. It should be noted that thermolysin cleavage removes the epitope for most tested glycan cap antibodies that showed very low specific binding to Jurkat-EBOV $GP_{CL}$.

| Antibody | Subject ID | VL | JL | VH | D | JH | CDRH3 length (amino acids) | Heavy chain VH mutations (%)* | Light chain VL mutations (%)** |
|---|---|---|---|---|---|---|---|---|---|
| EBOV-293 | 859 | *IGKV3-20*01* | *IGKJ5*01* | *IGHV1-69*09* | *IGHD3-9*01* | *IGHJ4*02/4*03* | 21 | 24/122 (19.6) | 14/107 (13) |
| EBOV-296 | 859 | *IGLV1-44*01* | *IGLJ3*02* | *IGHV4-30-4*01* | *IGHD3-10*01* | *IGHJ6*02* | 21 | 13/131 (9.9) | 7/111 (6.3) |
| BDBV-329 | 3 | *IGKV3-20*01* | *IGKJ4*01* | *IGHV1-69*10* | *IGHD1-26* | *IGHJ6*03* | 33 | 14/139 (10) | 4/108 (3.7) |
| BDBV-289 | 7 | *IGLV3-19*01* | *IGLJ2*01* | *IGHV1-69*06* | *IGHD5-24* | *IGHJ6*03* | 21 | 15/129 (11.6) | 4/108 (3.7) |
| BDBV-43 | 7 | *IGKV3-15*01* | *IGKJ5*01* | *IGHV1-69*09* | *IGHD6-6* | *IGHJ5*01* | 15 | 18/123 (14.6) | 7/106 (6.6) |
| EBOV-237 | EVD5 | *IGKV4-1*01* | *IGKJ2*01* | *IGHV4-4*02* | *IGHD3-3*01* | *IGHJ4*02* | 25 | 7/134 (5.2) | 5/102 (4.9) |
| EBOV-437 | 963 | *IGKV3-20*01* | *IGKJ2*02* | *IGHV1-69*06* | *IGHD3-10*01* | *IGHJ6*03* | 24 | 12/132 (9) | 5/108 (4) |
| EBOV-442 | 963 | *IGKV3-20*01* | *IGKJ1*01* | *IGHV3-15*01* | *IGHD3-16*01* | *IGHJ6*03* | 22 | 21/136 (15) | 7/108 (6.4) |
| EBOV-548 | 963 | *IGKV1-12*01* | *IGKJ2*01* | *IGHV3-30*02* | *IGHD6-19*01* | *IGHJ6*02* | 22 | 16/131 (12) | 10/108 (9.2) |

**Table S1. Summary of antibody germline characteristics, related to Table 1.**

*Heavy chain number of mutations in VH-encoded region from germline per total amino acids (%); **Light chain number of mutations in VL from germline per total amino acids (%)

| Map | EBOV GPΔMuc:BDBV289 | BDBV GPΔMuc:BDBV43 + ADI-15878 | EBOV GPΔMuc:EBOV-293 + EBOV-515 |
|---|---|---|---|
| **Sample vitrification** | - | - | - |
| Concentration | 3.5 mg/mL | 3 mg/mL | 3.5 mg/mL |
| Grid type | 1.2/1.3 C-Flat | 1.2/1.3 C-Flat | 1.2/1.3 Quantifoil |
| Detergent | 0.03 mM DDM | 0.01% (w/v) A8-35 | 0.01% (w/v) F-OM |
| Blot time | 3.5 s | 6 s | 6 s |
| **Data collection** | 18jan23c | 17sep13b | 19aug29a |
| Microscope | Titan Krios | Titan Krios | Talos Arctica |
| Voltage (kV) | 300 | 300 | 200 |
| Detector | K2 Summit | K2 Summit | K2 Summit |
| Recording mode | Counting | Counting | Counting |
| Magnification | 48,543 | 48,543 | 43,478 |
| Movie micrograph pixel size (Å) | 1.03 | 1.03 | 1.15 |
| Dose rate (e-/[(camera pixel)*s]) | 7.5 | 7.5 | 6.25 |
| Number of frames per movie | 32 | 32 | 50 |
| Frame exposure time (ms) | 250 | 250 | 250 |
| Movie micrograph exposure time (s) | 8 | 7.5 | 10.5 |
| Total dose (e-/Å$^2$) | 60 | 60 | 50 |
| Defocus range (µm) | -1 to -3.5 | -0.9 to -2.5 | 1 to -3.5 |
| **EM data processing** | - | - | - |
| Software used | Relion 3.1b | CryoSPARC 2.0 | CryoSPARC 2.0 |
| Number of movie micrographs | 1,061 | 2,753 | 796 |
| Number of particles in map | 32,108 | 383,783 | 79,722 |
| Symmetry | C3 | C3 | C3 |
| Map resolution (FSC 0.143;Å) | 3.8 | 4.1 | 4.25 |
| Map sharpening B-factor (Å$^2$) | -137 | -221 | -216 |
| **Structure building and validation** | - | - | - |
| Number of atoms in deposited model | 13,790 | 13,173 | 18,309 |
| GP1 | 5,682 | 5,628 | 5,352 |
| GP2 | 2,682 | 2,529 | 2,343 |
| BDBV289 | 5,426 | - | - |
| BDBV43 | - | 5,361 | - |
| EBOV-293 | - | - | 5,211 |
| EBOV-296 | - | - | - |
| EBOV-437 | - | - | - |
| EBOV-442 | - | - | - |
| BDBV-329 | - | - | - |
| BDBV-91 | - | - | - |
| glycans | 309 | 309 | 351 |
| MolProbity score | 0.62 | 0.92 | 1.07 |
| Clashscore | 0.33 | 0.54 | 0.97 |
| EMRinger score | 4.46 | 1.04 | 1.06 |
| RMSD from ideal | - | - | - |
| Bond length (Å) | 0.02 | 0.021 | 0.021 |
| Bond angle (degrees) | 1.65 | 1.81 | 1.82 |
| Ramachandran plot | - | - | - |
| Favored (%) | 99.65 | 96.47 | 95.91 |
| Allowed (%) | 0.35 | 3.16 | 3.56 |
| Outliers (%) | 0 | 0.37 | 0.53 |
| Average B-factor | 61 | 160 | 191 |

| Map | EBOV GPΔMuc:EBOV-296 + EBOV-515 | EBOV GPΔMuc:EBOV-437 + EBOV-515 | EBOV GPΔMuc:EBOV-442 + EBOV-515 |
|---|---|---|---|
| **Sample vitrification** | - | - | - |
| Concentration | 3.5 mg/mL | 3 mg/mL | 3 mg/mL |
| Grid type | 1.2/1.3 Quantifoil | 1.2/1.3 Quantifoil | 1.2/1.3 Quantifoil |
| Detergent | 0.01% (w/v) F-OM | 0.01% (w/v) F-OM | 0.01% (w/v) F-OM |
| Blot time | 6 s | 6 s | 6 s |
| **Data collection** | 19aug28b | 19aug24a | 19aug23b |
| Microscope | Titon Krios | Titan Krios | Titan Krios |
| Voltage (kV) | 300 | 300 | 300 |
| Detector | K2 Summit | K2 Summit | K2 Summit |
| Recording mode | Counting | Counting | Counting |
| Magnification | 48,543 | 48,543 | 48,543 |
| Movie micrograph pixel size (Å) | 1.03 | 1.03 | 1.03 |
| Dose rate (e-/[(camera pixel)*s]) | 6.43 | 6.43 | 6.43 |
| Number of frames per movie | 42 | 42 | 42 |
| Frame exposure time (ms) | 250 | 250 | 250 |
| Movie micrograph exposure time (s) | 10.5 | 10.5 | 10.5 |
| Total dose (e-/Å$^2$) | 50 | 50 | 50 |
| Defocus range (μm) | -1 to -3.5 | -1 to -3.5 | -1 to -3.5 |
| **EM data processing** | - | - | - |
| Software used | CryoSPARC 2.0 | Relion 3.1b | Relion 3.1b |
| Number of movie micrographs | 785 | 1,521 | 1,020 |
| Number of particles in map | 69,564 | 42,770 | 21,285 |
| Symmetry | C3 | C3 | C3 |
| Map resolution (FSC 0.143;Å) | 4.4 | 3.8 | 3.3 |
| Map sharpening B-factor (Å$^2$) | -126 | -142 | -92 |
| **Structure building and validation** | - | - | - |
| Number of atoms in deposited model | 18,570 | 18,981 | 18,783 |
| GP1 | 5,298 | 5,589 | 5,541 |
| GP2 | 2,388 | 2.454 | 2,373 |
| BDBV289 | - | - | - |
| BDBV43 | - | - | - |
| EBOV-293 | - | - | - |
| EBOV-296 | 5,481 | - | - |
| EBOV-437 | - | 5,535 | - |
| EBOV-442 | - | - | 5,571 |
| BDBV-329 | - | - | - |
| BDBV-91 | - | - | - |
| glycans | 309 | 393 | 534 |
| MolProbity score | 0.96 | 0.76 | 1.02 |
| Clashscore | 0.71 | 0.35 | 1.84 |
| EMRinger score | 0.93 | 3.44 | 3.62 |
| RMSD from ideal | - | - | - |
| Bond length (Å) | 0.021 | 0.021 | 0.021 |
| Bond angle (degrees) | 1.81 | 1.71 | 1.8 |
| Ramachandran plot | - | - | - |
| Favored (%) | 96.45 | 97.32 | 97.65 |
| Allowed (%) | 3.42 | 2.17 | 1.69 |
| Outliers (%) | 0.13 | 0.51 | 0.65 |
| Average B-factor | 201 | 145 | 128 |

| Map | BDBV GPΔMuc:BDBV329 + EBOV-515 | EBOV GPΔMuc:EBOV-237 + EBOV-515 |
|---|---|---|
| **Sample vitrification** | - | - |
| Concentration | 4 mg/mL | 3.1 mg/mL |
| Grid type | 1.2/1.3 Quantifoil | 1.2/1.3 Quantifoil |
| Detergent | 0.06 mM DDM | 0.06 mM |
| Blot time | 4.5 s | 4.5 s |
| **Data collection** | 19nov11g | 19nov12i |
| Microscope | Talos Arctica | Talos Arctica |
| Voltage (kV) | 200 | 200 |
| Detector | K2 Summit | K2 Summit |
| Recording mode | Counting | Counting |
| Magnification | 43,478 | 43,478 |
| Movie micrograph pixel size (Å) | 1.15 | 1.15 |
| Dose rate (e-/[(camera pixel)*s]) | 6.25 | 6.25 |
| Number of frames per movie | 42 | 42 |
| Frame exposure time (ms) | 250 | 250 |
| Movie micrograph exposure time (s) | 10.5 | 10.5 |
| Total dose (e-/Å$^2$) | 50 | 50 |
| Defocus range (µm) | -1 to -3.5 | -1 to -3.5 |
| **EM data processing** | - | - |
| Software used | Relion 3.1b | Relion 3.1b |
| Number of movie micrographs | 1,090 | 844 |
| Number of particles in map | 45,640 | 17,244 |
| Symmetry | C1 | C1 |
| Map resolution (FSC 0.143;Å) | 6.6 | 9.2 |
| Map sharpening B-factor (Å$^2$) | -242 | -369 |
| **Structure building and validation** | - | - |
| Number of atoms in deposited model | 16,721 | N/A |
| GP1 | 5,430 | N/A |
| GP2 | 2,118 | N/A |
| BDBV289 | - | - |
| BDBV43 | - | - |
| EBOV-293 | - | - |
| EBOV-296 | - | - |
| EBOV-437 | - | - |
| EBOV-442 | - | - |
| BDBV-329 | 3,770 | - |
| BDBV-91 | - | - |
| glycans | 267 | N/A |
| MolProbity score | 0.9 | N/A |
| Clashscore | 0.4 | N/A |
| EMRinger score | 0.98 | N/A |
| RMSD from ideal | - | - |
| Bond length (Å) | 0.022 | N/A |
| Bond angle (degrees) | 1.912 | N/A |
| Ramachandran plot | - | - |
| Favored (%) | 96.19 | N/A |
| Allowed (%) | 3.24 | N/A |
| Outliers (%) | 0.56 | N/A |
| Average B-factor | 236 | N/A |

**Table S2. Cryo-EM data collection and validation statistics, related to Figure 2 and Figure 3.**

| Residue number | Total Contacts | Residue | Number of amino acid contacts with GP | | | | | | | | |
|---|---|---|---|---|---|---|---|---|---|---|---|
| | | | EBOV-548 | EBOV-437 | 13C6 | BDBV-329 | EBOV-442 | BDBV-43 | BDBV289 | EBOV-296 | EBOV-293 |
| 116 | 13 | P | 1 | 3 | 4 | 5 | | | | | |
| 117 | 5 | D | 3 | | 1 | 1 | | | | | |
| 118 | 5 | G | | | | 5 | | | | | |
| 144 | 7 | T | | | 1 | 6 | | | | | |
| 221 | 4 | Q | | 3 | | | | | 1 | | |
| 222 | 0 | A | | | | | | | | | |
| 223 | 15 | T | | 4 | 3 | 2 | 3 | | 3 | | |
| 227 | 1 | T | | | | 1 | | | | | |
| 229 | 18 | E | 2 | 2 | | 7 | | | | 7 | |
| 231 | 26 | E | 7 | 4 | | 2 | 8 | | 3 | 2 | |
| 233 | 13 | L | | 2 | 1 | | 2 | | 8 | | |
| 239 | 5 | L | | | | | | 5 | | | |
| 240 | 11 | T | | | 4 | | | 7 | | | |
| 241 | 8 | Y | | 3 | | 1 | | | 4 | | |
| 242 | 4 | V | | 2 | | | 1 | 1 | | | |
| 243 | 10 | Q | | 4 | | | 3 | | 2 | 1 | |
| 252 | 4 | F | | | | | | | | 4 | |
| 256 | 25 | L | 1 | 6 | | | 5 | 5 | 3 | 1 | 4 |
| 259 | 9 | T | | 2 | | | | | 4 | 2 | 1 |
| 260 | 12 | I | 2 | | | | 1 | 2 | 2 | 1 | 4 |
| 262 | 1 | T | | | | | | 1 | | | |
| 263 | 15 | S | | | | 4 | | 8 | 3 | | |
| 264 | 1 | G | | | | 1 | | | | | |
| 265 | 28 | K | 4 | 4 | | | | | 11 | 3 | 6 |
| 267 | 8 | S | | | | | | 2 | | | 6 |
| 268 | 32 | N | | | | 1 | 1 | | | 18 | 12 |
| 269 | 26 | T | | 1 | 3 | 3 | 2 | 4 | 4 | 5 | 4 |
| 270 | 49 | T | | 2 | 19 | 2 | 4 | 10 | 10 | 1 | 1 |
| 271 | 31 | G | 2 | 5 | 3 | 3 | 3 | 3 | 3 | 3 | 6 |
| 272 | 65 | K | 8 | 14 | 10 | 7 | 5 | | 16 | 5 | |
| 273 | 67 | L | 10 | 1 | 10 | 12 | 11 | 8 | 3 | 9 | 3 |
| 274 | 46 | I | 8 | 10 | 5 | 2 | 9 | | 6 | 5 | 1 |
| 275 | 163 | W | 6 | 26 | | 3 | 14 | 29 | 10 | 54 | 21 |
| 276 | 46 | K | 5 | 3 | | 1 | 3 | 6 | 6 | 15 | 7 |
| 277 | 15 | V | | 5 | | | 4 | 2 | 1 | 1 | 2 |
| 278 | 29 | N | 5 | 2 | | | 3 | 3 | 4 | 6 | 6 |
| 280 | 14 | E | | | | | | | 6 | 8 | |
| 281 | 6 | I | 2 | 2 | | | | | 2 | | |
| 282 | 4 | D | | | | | | | 4 | | |

**Table S3. Number of glycan cap antibody contacts with GP, related to Figure 3.**

Glycan cap antibody contacts on EBOV/BDBV GP determined by UCSF Chimera software with a default cutoff value of -0.4 Å and an allowance of 0 Å. EBOV-548 from PDB 6UYE, 13C6 from PDB 5KEL. Gradient is green-to-red, with green being fewer contacts and red being the most contacts.

| | |
|---|---|
| **Data Collection** | |
| Resolution[a] (Å) | 50.00 - 3.00 (3.05 - 3.00) |
| Space group | $P6_1$ |
| Unit cell (Å)<br>(°) | 94.46 94.46 94.911<br>90 90 120 |
| Total Reflections[a]<br><br>Unique Reflections[a]<br><br>Reflections used in Refinement[a] | 1103684<br><br>9472<br><br>9473 |
| Multiplicity[a] | 4.7 (4.5) |
| Completeness[a] (%) | 97.8 (99.2) |
| I/σ(I)[a] | 8.6 (1.4) |
| $R_{meas}$[a,b] | 0.19 (1.38) |
| $R_{pim}$[a,c] | 0.083 (0.612) |
| CC 1/2[d] | -0.295 |
| Wilson B ($Å^2$) | 68.35 |
| **Refinement** | |
| $R_{work}$[e] (%)<br>$R_{free}$[f] (%) | 0.22<br>0.27 |
| RMSD[g] (bonds) (Å) | 0.002 |
| RMSD[g] (angles) (°) | 0.51 |
| Ramachandran favored (%)<br>Ramachandran outliers (%) | 94.8<br>0 |
| Average B-factor ($Å^2$)<br>Macromolecules<br>PEG | 85.4<br>85.5<br>53.9 |

**Table S4. Crystallographic Data Collection and Refinement Statistics for BDBV-289 Fab, related to Figure 4.**

[a] Values in parentheses are for the highest-resolution shell.
[b] $R_{meas} = \Sigma_{hkl}\{N(hkl)/[N(hkl) - 1]\}^{1/2} \times \Sigma_i|I_i(hkl) - \langle I(hkl)\rangle| / \Sigma_{hkl} \Sigma_i I_i(hkl)$
[c] $R_{pim} = \Sigma_{hkl}\{1/[N(hkl) - 1]\}^{1/2} \times \Sigma_i|I_i(hkl) - \langle I(hkl)\rangle| / \Sigma_{hkl} \Sigma_i I_i(hkl)$
[d] $CC_{1/2} = \Sigma(x - \langle x\rangle)(y - \langle x\rangle)/[\Sigma(x - \langle x\rangle)^2\Sigma(y - \langle y\rangle)^2]^{1/2}$
[e] $R_{work} = (\Sigma_{hkl}||F_{obs}| - k\,|F_{calc}||)/(\Sigma_{hkl}\,|F_{obs}|)$.
[f] $R_{free}$ is the same as $R_{work}$ with 5% of reflections chosen at random and omitted from refinement.
[g] RMSD, root mean square deviation.
